# Arrayed CRISPR/Cas9 Loss-Of-Function Screen in a Neuronal Model of Adaptor Protein Complex 4 Deficiency Identifies Modulators of ATG9A Trafficking

**DOI:** 10.1101/2025.02.22.639634

**Authors:** Marvin Ziegler, Cedric Böger, Julian E. Alecu, Hyo-Min Kim, Afshin Saffari, Alexandra K. Davies, Mustafa Sahin, Darius Ebrahimi-Fakhari

## Abstract

Biallelic loss-of-function variants in the adaptor protein complex 4 (AP-4) disrupt trafficking of transmembrane proteins at the *trans*-Golgi network, including the autophagy-related protein 9A (ATG9A), leading to childhood-onset hereditary spastic paraplegia (AP-4-HSP). AP-4-HSP is characterized by features of both a neurodevelopmental and degenerative neurological disease. To investigate the molecular mechanisms underlying AP-4-HSP and identify potential therapeutic targets, we conducted an arrayed CRISPR/Cas9 loss-of-function screen of 8,478 genes, targeting the ‘druggable genome’, in a human neuronal model of AP-4 deficiency. Through this phenotypic screen and subsequent experiments, key modulators of ATG9A trafficking were identified, and complementary pathway analyses provided insights into the regulatory landscape of ATG9A transport. Knockdown of *ANPEP* and *NPM1* enhanced ATG9A availability outside the *trans*-Golgi network, suggesting they regulate ATG9A localization. These findings deepen our understanding of ATG9A trafficking in the context of AP-4 deficiency and offer a framework for the development of targeted interventions for AP-4-HSP.

## INTRODUCTION

Approximately one-third of all genes in the eukaryotic genome are predicted to encode transmembrane proteins that must traffic through various cellular compartments to function effectively.^1,2^ Defective transport of these proteins has been increasingly linked to a range of neurological diseases, including frontotemporal dementia, Charcot-Marie-Tooth disease, and several types of hereditary spastic paraplegia (HSP).^3,4^ Among these, adaptor protein complex 4 (AP-4)-associated hereditary spastic paraplegia (AP-4-HSP) has emerged as a prototypical disorder of impaired protein trafficking. Clinically, AP-4-HSP manifests in early childhood with developmental delay, intellectual disability, progressive spasticity leading to loss of ambulation, epilepsy, secondary microcephaly, and developmental brain malformations.^5–8^ The condition arises from biallelic loss-of-function variants in one of four genes (*AP4B1*, *AP4E1*, *AP4M1*, *AP4S1*) encoding the subunits of the heterotetrameric AP-4 complex, which is critical for sorting cargo proteins from the *trans*-Golgi network (TGN) to early and late endosomes and pre-autophagosomal structures (PAS).^5,6^

Recent studies have identified ATG9A, the sole transmembrane protein in the autophagic machinery, as an important cargo of AP-4.^9–11^ In AP-4-deficient cells, ATG9A accumulates in the TGN and is depleted from peripheral compartments. In neurons, this mislocalization leads to deficits in autophagy, impaired axon maintenance, and length-dependent axonal degeneration.^11–16^ However, residual autophagic activity and the presence of ATG9A outside the TGN suggest an alternative, AP-4-independent export mechanism.^9,11^ This raises critical questions about how ATG9A trafficking is regulated in the context of AP-4 deficiency.

Genome-wide functional genomics screens have emerged as powerful tools to uncover regulators of cellular signaling. However, traditional approaches such as loss-of-function screening using small interfering RNA (siRNA) libraries are often limited by poor reproducibility due to inconsistent gene knockdown and off-target effects.^17,18^ Advances in CRISPR/Cas9-based gene interference, particularly its application in high-throughput screening assays, have addressed many of these limitations, ushering in a new era in functional genomics.^19^ While several large-scale CRISPR screens have been successfully conducted in recent years - primarily as pooled screens where cells are bulk transfected with a single guide RNA (sgRNA) library, selected for specific phenotypes, and analyzed using next-generation sequencing - these approaches pose significant challenges for assessing complex cellular phenotypes.^20,21^

In contrast, arrayed CRISPR screens offer a microplate-based format in which gene modifications are performed individually in each well. This format eliminates the need for complex data deconvolution, facilitates the analysis of intricate readouts such as high-content microscopy, and leverages established platforms for siRNA and small molecule screening.^19^ Consequently, arrayed screens provide a robust and scalable approach for functional genomics with broad applications, including drug target discovery.^22^ However, only a limited number of arrayed CRISPR screens have been conducted to date.^23–25^

Here, we present a large-scale CRISPR/Cas9 loss-of-function screen designed to identify modulators of ATG9A trafficking in a neuronal model of AP-4 deficiency, specifically targeting the ‘druggable genome’. By combining high-content imaging-based multiparametric analysis with a tailored pathway analysis for arrayed screening, we identify novel genes and pathways that modulate ATG9A trafficking. These findings reveal promising targets for drug development and open new avenues for therapeutic interventions in AP-4-HSP.

## RESULTS

### Establishing a Phenotypic High-Content Screen for Modulators of ATG9A Trafficking

We previously developed and validated a high-content imaging-based assay to measure intracellular ATG9A distribution as a surrogate for AP-4 function.^26,27^ To identify modulators of ATG9A trafficking in AP-4-deficient cells, we have now leveraged this assay to conduct an arrayed CRISPR/Cas9-based loss-of-function screen (*Fig. 1A*). Given that AP-4-HSP predominantly affects cortical and spinal neurons, we utilized the SH-SY5Y neuroblastoma cell line, which can be differentiated with retinoic acid into a neuron-like state resembling cholinergic neurons.^28^ For this study, we employed SH-SY5Y cells stably expressing wild-type Cas9, with and without expression of a sgRNA to knockout *AP4B1.*^10^ Western blot analysis confirmed Cas9 expression and the absence of AP4B1 protein in *AP4B1^KO^* cells, which also displayed reduced levels of AP4E1, consistent with diminished overall AP-4 complex formation. As previously observed in other AP-4-deficient cells,^9,10,14^ ATG9A protein levels were elevated in *AP4B1^KO^*cells (*Fig. 1B, S1)*.

**Figure 1:**
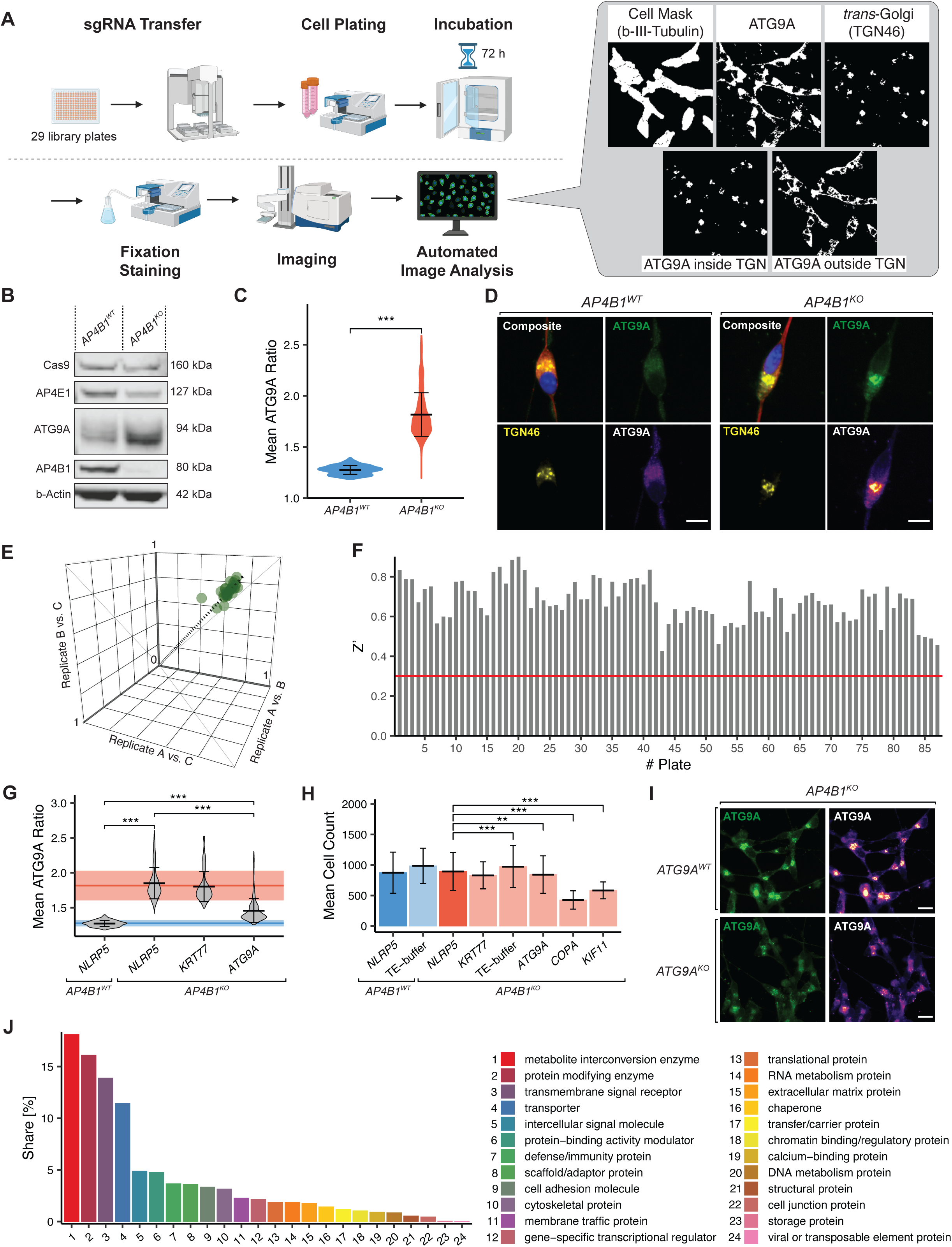
Establishment of a High-Content CRISPR/Cas9 Loss-Of-Function Screening Platform. *A)* Experimental workflow for CRISPR/Cas9 screen. Control and library sgRNAs were transferred into 384-well plates preloaded with Lipofectamine™ RNAiMAX. Differentiated Cas9 expressing SH-SY5Y cells were plated on top and incubated with sgRNAs for 72 hours. Cells were subsequently fixed and stained with Hoechst 33258 (nuclear marker), β-Tubulin III (cytoplasmic marker), TGN46 (*trans*-Golgi network marker), and ATG9A (ATG9A compartment marker). Plates were imaged using an automated high-content imaging platform (Molecular Devices ImageXpress Micro Confocal Laser System and PAA GX™ robot arm), and images were analyzed using an automated pipeline in MetaXpress to segment cells and calculate the proportion of ATG9A signal overlapping with the TGN. This figure was partially created with BioRender.com. Cellular compartment masks derived from MetaXpress. *B)* Western blot analysis of whole-cell lysates from differentiated *AP4B1^KO^* and *AP4B1^WT^* SH-SY5Y cells. *AP4B1^KO^* cells showed loss of AP4B1 protein and reduced levels of AP4E1, indicating successful *AP4B1* knockout and decreased AP-4 complex formation. ATG9A levels were increased in *AP4B1^KO^* cells. Both cell lines demonstrated stable expression of Cas9. *C)* ATG9A ratio of untransfected *AP4B1^KO^* and *AP4B1^WT^* cells, based on all wells from the primary screen. *AP4B1^KO^* cells showed a significant increase in the ATG9A ratio, consistent with accumulation of ATG9A in the TGN. Statistical analysis: Two-sided Student’s t-test, p < 0.001 (***), Cohen’s d ≈ 2.74. *D)* Representative images of *AP4B1^KO^* and *AP4B1^WT^* SH-SY5Y cells stained with antibodies against ATG9A and TGN46. *AP4B1^KO^* cells exhibited perinuclear accumulation of ATG9A overlapping with the TGN. Pseudocolored images depict grayscale intensity. Scale bar = 10 µm. *E)* Pearson correlation coefficients between replicates across all plates in the primary screen. High correlation between replicates demonstrated low interplate variability. Mean Pearson correlation coefficient: 0.907 ± 0.035. *F)* Z’ robust for all 87 screened plates confirmed reliable assay performance. Horizontal line indicates predefined quality control threshold (Z’ > 0.3). *G)* ATG9A Ratio for *AP4B1^KO^* and *AP4B1^WT^* cells transfected with sgRNAs targeting non-essential genes (*NLRP5* and *KRT77*), *ATG9A*, and essential genes (*COPA* and *KIF11*). Red lines represent the ATG9A ratio ± SD for untransfected *AP4B1^KO^* cells (negative control), and blue lines represent the same for *AP4B1^WT^* cells (positive control). Statistical analysis: One-way ANOVA, F(5) = 1908, p < 0.001, η² = 0.58. Tukey HSD post hoc test revealed significant differences between groups p < 0.001 (***). *H)* Mean cell counts for *AP4B1^KO^* (red) and *AP4B1^WT^* (blue) cells, including untransfected controls (TE-buffer) and cells transfected with sgRNAs targeting non-essential (*NLRP5*, *KRT77*) and essential (*COPA*, *KIF11*) genes, as well as ATG9A. Transfection with sgRNAs targeting essential genes significantly reduced cell counts, confirming efficient transfection and gene knockout. Statistical analysis: One-way ANOVA, F(7) = 163, p < 0.001, η² = 0.13. Dunnett’s post hoc test identified significant group differences p < 0.01 (**) and p < 0.001 (***). *I)* Representative images of *AP4B1^KO^*/*ATG9A^WT^* (top panel) and *AP4B1^KO^*/*ATG9A^KO^* (bottom panel) SH-SY5Y cells stained for ATG9A and TGN46. Pseudocolored images depict grayscale intensity. Scale bar = 20 µm. *J)* Frequency of Panther GO terms associated with genes in the ‘Synthego Human Druggable Genome’ library.

For the CRISPR/Cas9-based loss-of-function screen, SH-SY5Y cells were reversely transfected with the ‘Synthego Human Druggable Genome’ sgRNA library and incubated for 72 hours. To ensure high consistency and minimize variability, we used automated systems, including sgRNA transfer, pipetting, and liquid dispensing. High-content imaging at 40× magnification was performed using the IXM high-content imaging system, followed by automated analysis of individual cells with hundreds of cells per well (*AP4B1^WT^*: 930.342 ± 318.332, *AP4B1^KO^*: 845.354 ± 289.564) using a custom pipeline developed with MetaXpress software. Cellular compartments were labeled with four markers: Hoechst 33258 (nucleus), β-Tubulin III (cytoskeleton), TGN46 (*trans*-Golgi network), and ATG9A (ATG9A compartment). To quantify ATG9A distribution, we calculated the ratio of ATG9A localized inside versus outside the TGN. Consistent with previous findings,^9,10,26^ *AP4B1^KO^*cells showed distinct perinuclear accumulation of ATG9A overlapping with the TGN (*Fig. 1C, D*), resulting in a significantly elevated ATG9A Ratio (*AP4B1^WT^*: 1.276 ± 0.043, *AP4B1^KO^*: 1.817 ± 0.213; p < 0.001, Cohen’s d ≈ 2.74).

The ‘Synthego Human Druggable Genome’ sgRNA library targets 8,478 genes, arrayed in a 384-well format, with designated spaces for assay-specific controls. To address potential off-target effects and transfection toxicity, we included negative controls such as *AP4B1^KO^* cells transfected with sgRNAs targeting the non-essential genes *NLRP5* and *KRT77*, as well as untransfected *AP4B1^KO^* cells. Positive controls included *AP4B1^WT^* cells, either untransfected or transfected with sgRNAs targeting *NLRP5*. Additionally, *AP4B1^KO^* cells transfected with sgRNAs targeting essential genes *COPA* and *KIF11* served as controls to assess transfection efficiency. sgRNAs targeting *ATG9A* in *AP4B1^KO^* cells were used to evaluate the assay’s sensitivity to dynamic changes in ATG9A trafficking (*Fig. S2A*).

Each library plate was screened in three technical replicates with high consistency across replicates demonstrated by Pearson’s correlation coefficient (0.907 ± 0.035) (*Fig. 1E*). Quality metrics for high-throughput screening, including Z’ and SSMD (robust versions included), exceeded predefined thresholds (Z’ and Z’ robust > 0.3; SSMD and SSMD robust > 5) (*Fig. 1F*, *Fig. S2B–D*). Knockout of non-essential genes did not alter the ATG9A ratio in either *AP4B1^KO^* or *AP4B1^WT^* cells (*Fig. 1G*). However, minor reductions in cell count suggested low levels of toxicity from off-target effects (*Fig. 1H*). Knockout of *ATG9A* similarly reduced cell counts but significantly decreased the ATG9A ratio in *AP4B1^KO^* cells, nearly matching levels observed in *AP4B1^WT^* cells. This reduction was accompanied by diminished perinuclear ATG9A accumulation (*Fig. 1I*). As expected, knockout of essential genes induced apoptosis. For instance, sgRNAs targeting *COPA* reduced cell counts by approximately 52%, reflecting efficient transfection and knockout. Interestingly, *KIF11* knockout decreased cell counts by only 35%, possibly due to the reduced importance of *KIF11* in postmitotic SH-SY5Y cells (*Fig. 1H*).

Finally, analysis of the sgRNA library using Panther gene ontology (GO) terms revealed that most targeted genes encode metabolite interconversion enzymes, protein-modifying enzymes, transmembrane signal receptors and transporters (*Fig. 1J*).

### Primary and Secondary Screens Identify Key Modulators of ATG9A Trafficking

As anticipated, the majority of the 8,478 genes screened did not significantly impact ATG9A distribution, as measured by the mean z-score robust of the ATG9A ratio across all three replicates (*Fig. 2A, B*). However, 1,041 genes (12.28%) were found to decrease the cell count (z-score robust < −3 in two or more replicates) and were excluded from further analysis due to potential cytostatic or cytotoxic effects. Ultimately, 92 genes met the predefined threshold for ATG9A ratio reduction (z-score robust < −3 in all replicates), with a mean z-score robust across all replicates of −4.794 ± 1.123. This represents a hit rate of 1.09%, comparable to other high-throughput screening assays.^29^

**Figure 2.**
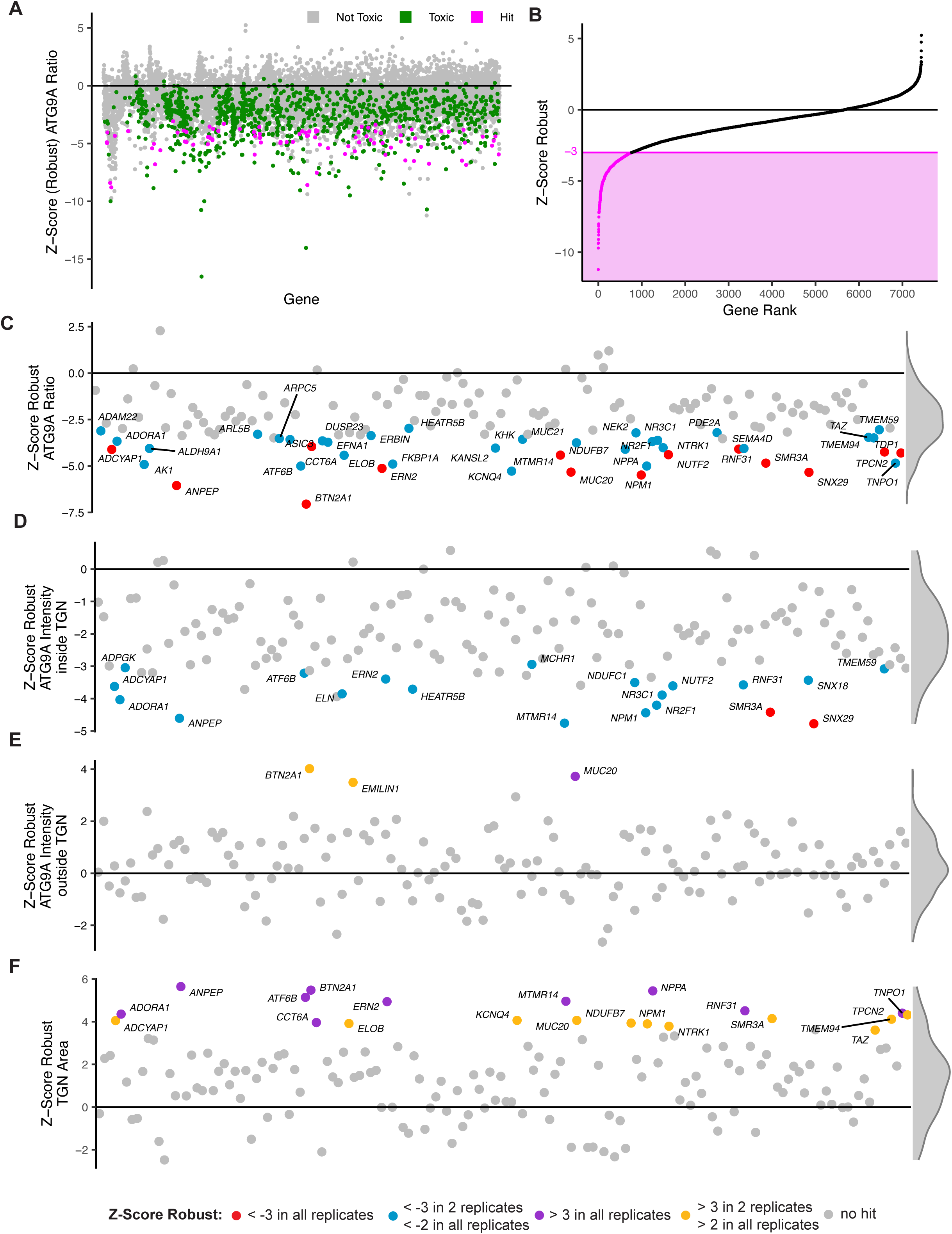
Primary and Secondary Screens Identify Modulators of ATG9A Trafficking. *A)* Distribution of robust z-scores from the primary screen. Genes are categorized as toxic (z-score robust for cell count < −3 in at least two replicates), not-toxic, or hits (not-toxic & z-score robust for ATG9A ratio < −3 in all replicates). A total of 92 genes were identified as hits, while 1,041 genes were classified as toxic. *B)* Ranked distribution of robust z-scores for all screened genes. Genes with a mean robust z-score < −3 across all replicates are highlighted. While only 92 genes met the stringent cut-off criteria of robust z-scores < −3 in all replicates and were classified as hits, 764 genes exhibited a mean robust z-score < −3 across all replicates, indicating a strong but less reproducible impact on ATG9A trafficking. *C)* Results of the secondary screen demonstrating robust z-scores for the ATG9A ratio. A total of 14 genes were confirmed as hits (robust z-scores < −3 in all replicates), additionally 30 genes met the less stringent criteria of robust z-scores < −3 in two replicates and < −2 in all replicates. *D)* Robust z-scores for ATG9A intensity inside the TGN in the secondary screen. Most hits reduced the ATG9A ratio by decreasing ATG9A intensity within the TGN. *E)* Robust z-scores for ATG9A intensity outside the TGN. Most genes had minimal impact on ATG9A intensity outside the TGN. *F)* Robust z-scores for TGN area. Several genes affected TGN morphology, as evidenced by an increase in TGN area, potentially indicating early toxicity or structural reorganization. Density distribution of robust z-scores displayed on the right in *C)* – *F)*.

To refine our findings and eliminate false positives, we conducted a secondary screen on the 92 identified hits. Logistical circumstances enabled us to rescreen up to 150 genes. Therefore, additional 58 genes were selected from the top of the list of genes meeting a slightly less stringent cutoff (ATG9A ratio: z-score robust < −3 in two replicates, z-score robust < −2 in all replicates, ranked by mean z-score robust of all replicates) (*Fig. S3A*). We followed the same experimental protocol as in the primary screen but extended the analysis to include a multi-parametric *trans*-Golgi network (TGN) morphology characterization, with the goal of identifying confounding effects on Golgi morphology that could impact the ATG9A ratio. Ultimately, 44 genes were confirmed to significantly impact ATG9A translocation: 14 genes consistently met the stringent z-score robust < −3 threshold across all replicates, while the remaining 30 genes satisfied a z-score robust < −2 in all replicates and < −3 in at least two replicates (*Fig. 2C*).

Most identified genes reduced the ATG9A ratio by decreasing ATG9A signal intensity inside the TGN, with minimal changes to ATG9A levels outside the TGN (*Fig. 2D*). Interestingly, three genes - *BTN2A1*, *EMILIN1*, and *MUC20* - primarily increased ATG9A signal intensity outside the TGN (*Fig. 2E*). Additionally, several genes were found to enlarge the TGN area, potentially indicating either a reorganization of the TGN or early signs of cytotoxicity (*Fig. 2F*). The exact biological significance of this TGN enlargement remains unclear and warrants further investigation.

The data from primary and secondary screens are provided as supplementary files (*Supplementary Files 1 and 2*), offering a comprehensive resource for exploring the genetic modulators of ATG9A trafficking.

### Pathway Analysis Illustrates Modulatory Landscape of ATG9A Trafficking

Hit identification in high-throughput screens based solely on ranked readout measures often overlooks the cumulative effects of functionally related genes that do not individually meet hit criteria. To address this limitation, we developed a customized pathway analysis specifically for arrayed high-throughput screens. Unlike conventional pathway analysis tools, our method incorporates the robust z-score for each gene rather than relying solely on its rank position. Inspired by an approach previously used in siRNA screens,^30^ we utilized a model that calculates the cumulative z-score robust adjusted for the pathway size (Pathway Impact (PI) Score) and determines the likelihood of obtaining a given PI score by chance (p-value). This model was applied to data from our primary screen. Ultimately, 463 out of 15,365 pathways (3.01%) were identified as significantly affected (*Fig. 3A*, *Supplementary File 4*).

**Figure 3:**
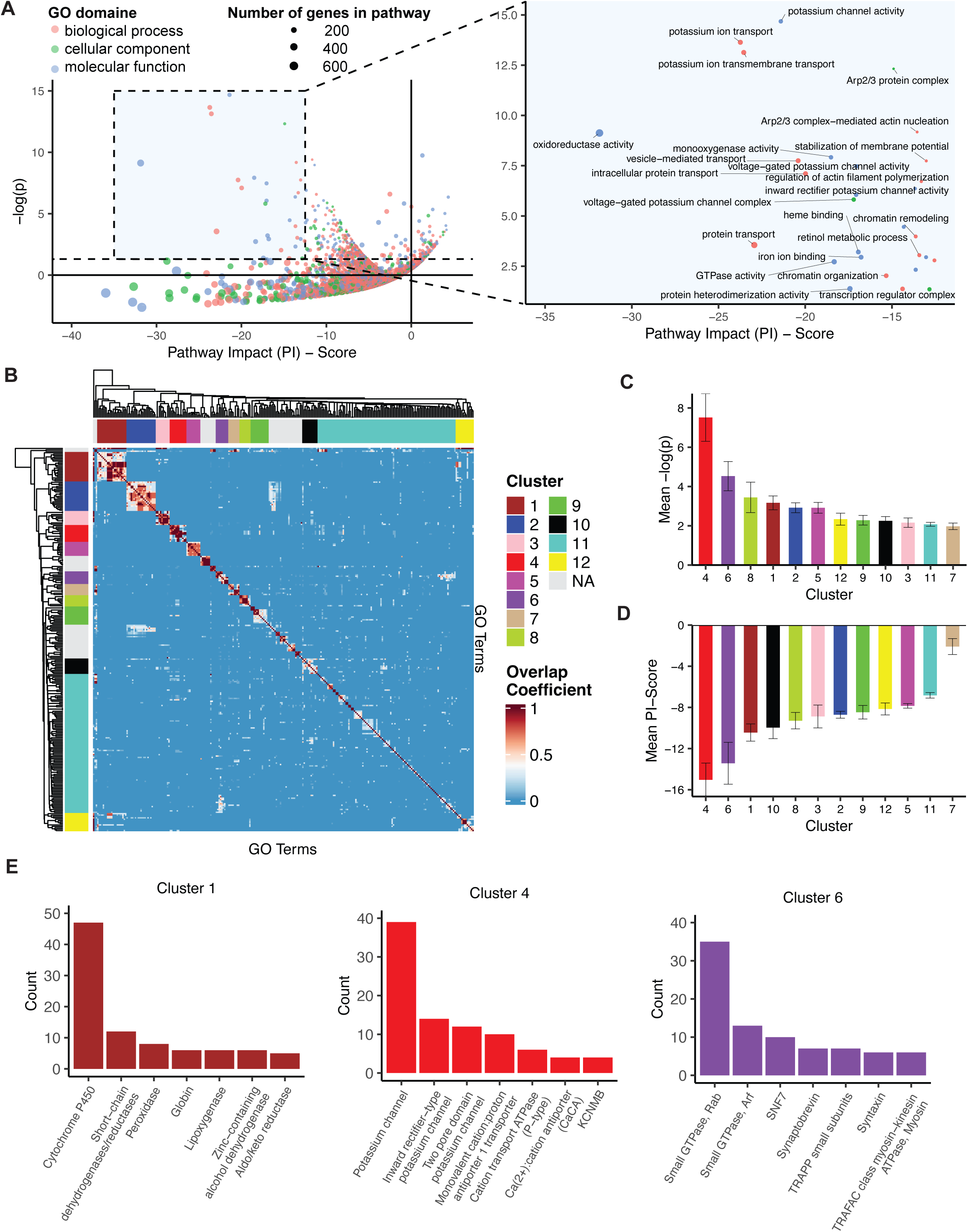
Pathway Analysis Reveals the Modulatory Landscape of ATG9A Trafficking. *A)* Pathway analysis of primary screen results. Each dot represents a GO term. Pathway Impact (PI) score reflects the cumulative robust z-score of all genes associated with the respective GO term adjusted for its size. Dot size corresponds to number of genes linked to the GO term; color indicates gene ontology branch. Detailed view of significant pathways with PI scores between −35 and −12.5 on the right, selected pathways labeled for clarity. *B)* Heatmap of genetic overlap between GO terms sorted by average linkage clustering. Overlap coefficients range from 0 (no genetic overlap) to 1 (complete genetic overlap of the smaller GO term with the larger GO term). Color bars on heatmap edges indicate clusters identified with the Dynamic Tree Cut algorithm. *C)* Mean p-values and standard deviations across pathways associated with identified clusters, highlighting their statistical significance. *D)* Mean PI scores and standard deviations across pathways associated with identified clusters, indicating cumulative impact of genes on ATG9A trafficking within each cluster. *E)* Protein families and number of proteins from each family associated with top three clusters: Cluster 1 (oxidative degradation, 192 proteins), cluster 4 (potassium channel activity, 133 proteins), and cluster 6 (vesicular transport, 308 proteins).

To better understand these pathways, we grouped functionally similar pathways into broader clusters by calculating genetic overlaps and constructing a similarity matrix. Following average linkage clustering, we applied the Dynamic Tree Cut algorithm – commonly used in gene expression analysis within the WGCNA package^31,32^ – resulting in 12 distinct clusters (*Fig. 3B*). Each cluster was manually annotated with umbrella terms reflecting its primary function (*Table 1*). For each cluster, we calculated the mean PI Score and mean z-score (≙ mean p-value) (*Fig. 3C, D*). Cluster 4, encompassing GO terms related to potassium channel activity, exhibited the lowest mean z-score and the highest mean PI Score, indicating a substantial impact on ATG9A trafficking. Cluster 1, associated with oxidative degradation, also strongly influenced ATG9A distribution. Notably, cluster 6, comprising GO terms related to vesicular transport, demonstrated significant effects on ATG9A redistribution. Conversely, cluster 11 contained 99 pathways with no meaningful connections and was excluded from further analysis.

**Table 1:**
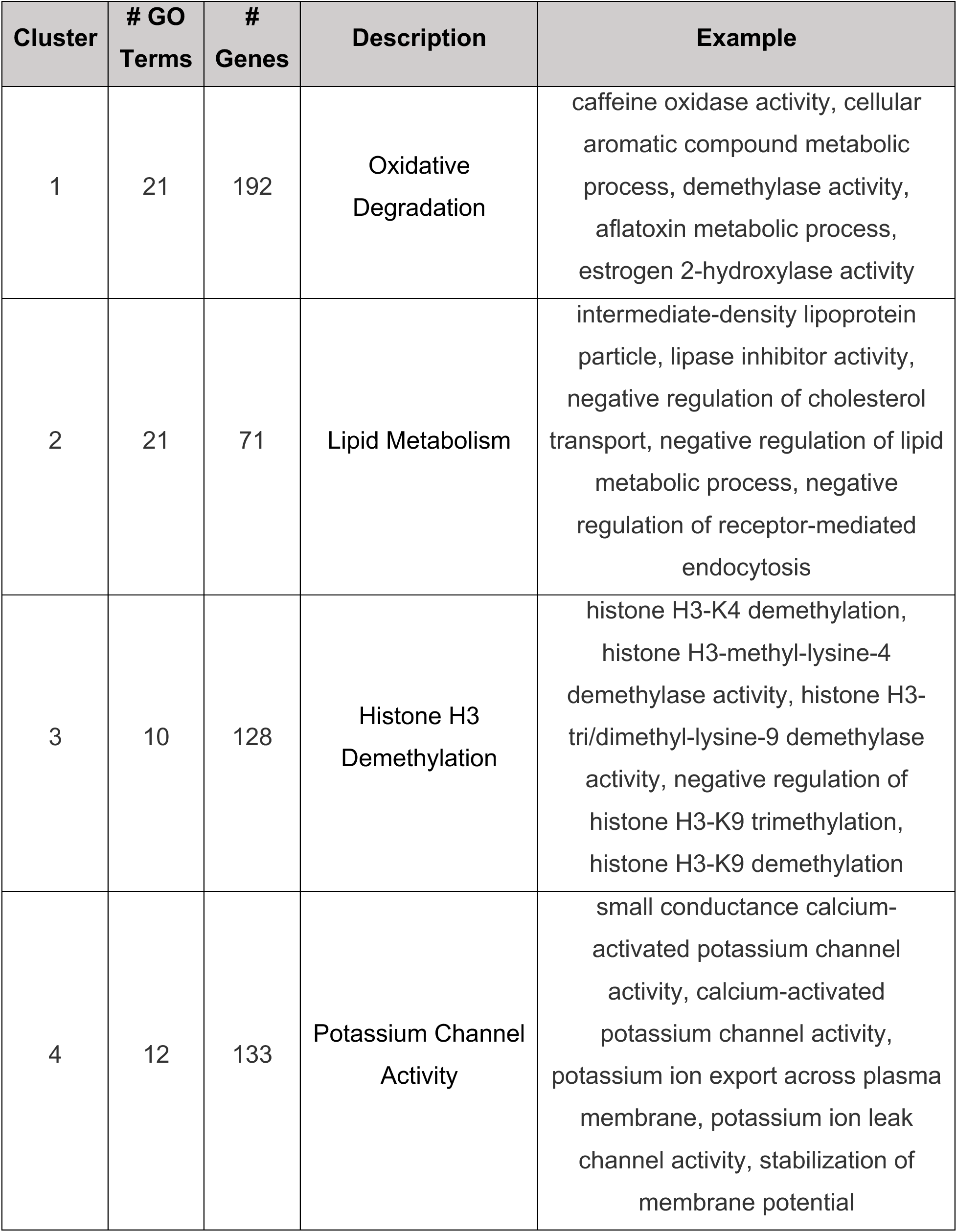
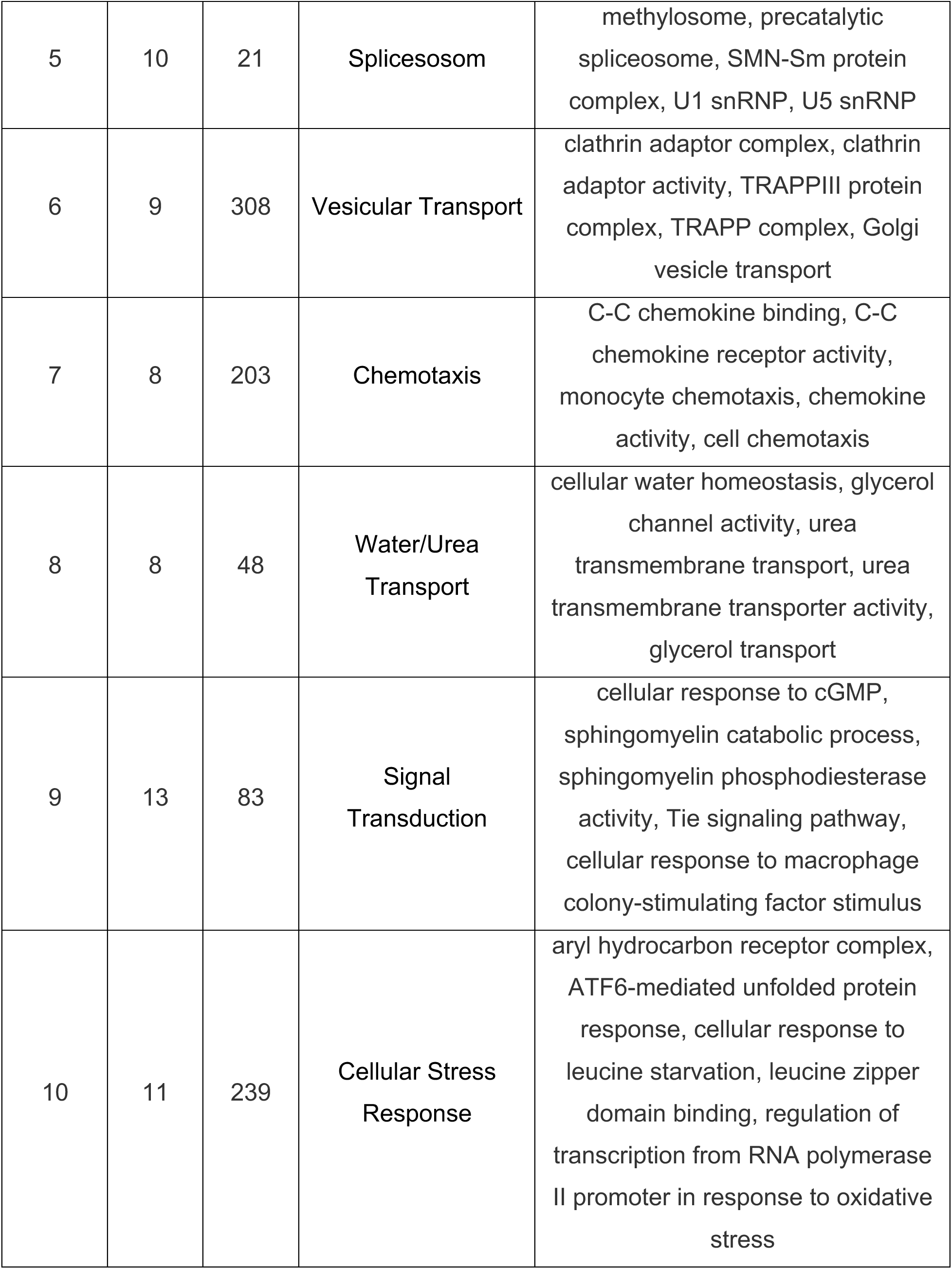
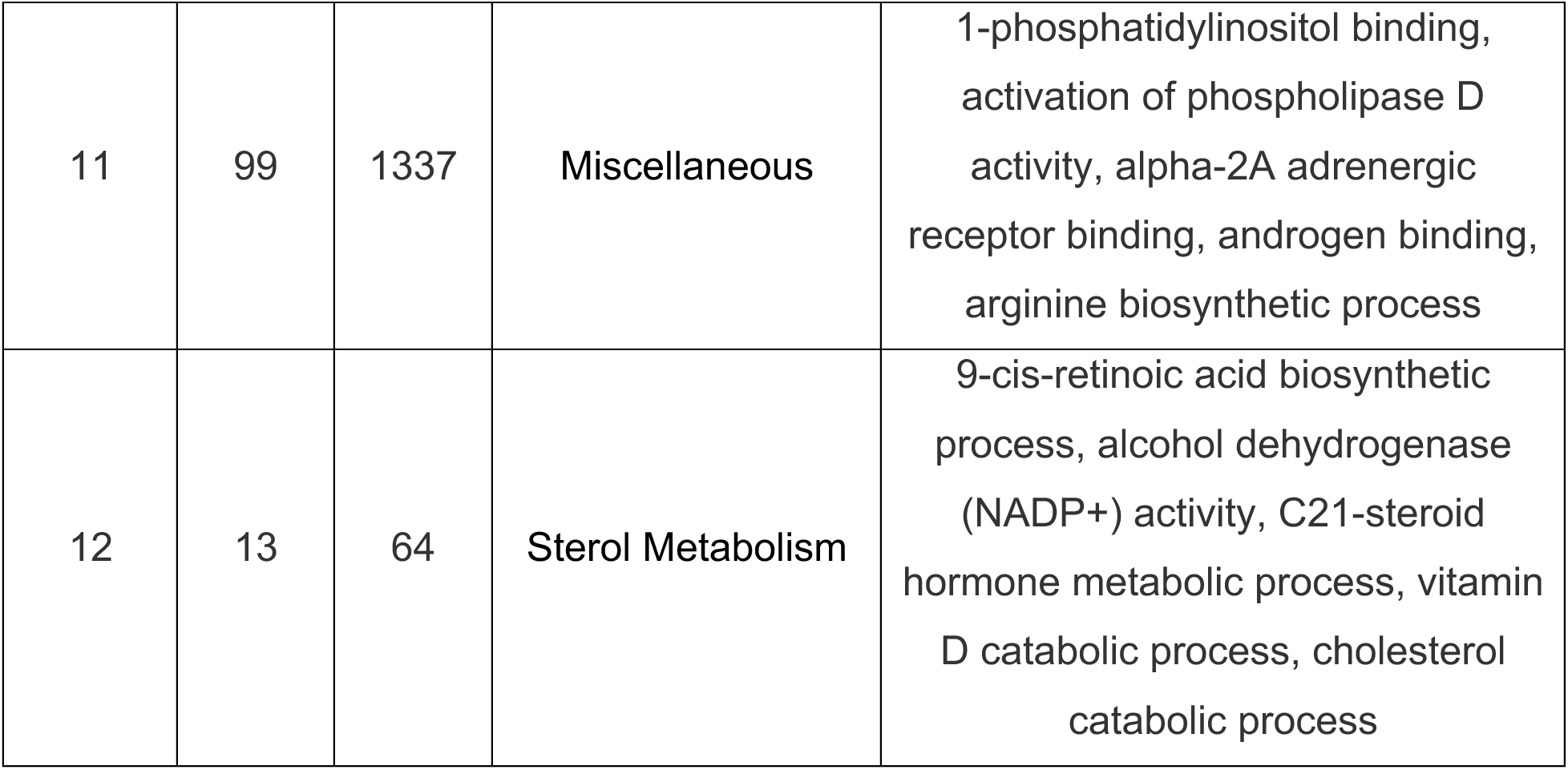
Pathway Clusters Identified by Average-Linkage Hierarchical Clustering Based On Genetic Overlap Between Significantly Affected Pathways.

| Cluster | # GO Terms | # Genes | Description | Example |
| --- | --- | --- | --- | --- |
| 1 | 21 | 192 | Oxidative Degradation | caffeine oxidase activity, cellular aromatic compound metabolic process, demethylase activity, aflatoxin metabolic process, estrogen 2-hydroxylase activity |
| 2 | 21 | 71 | Lipid Metabolism | intermediate-density lipoprotein particle, lipase inhibitor activity, negative regulation of cholesterol transport, negative regulation of lipid metabolic process, negative regulation of receptor-mediated endocytosis |
| 3 | 10 | 128 | Histone H3 Demethylation | histone H3-K4 demethylation, histone H3-methyl-lysine-4 demethylase activity, histone H3-tri/dimethyl-lysine-9 demethylase activity, negative regulation of histone H3-K9 trimethylation, histone H3-K9 demethylation |
| 4 | 12 | 133 | Potassium Channel Activity | small conductance calcium-activated potassium channel activity, calcium-activated potassium channel activity, potassium ion export across plasma membrane, potassium ion leak channel activity, stabilization of membrane potential |
| 5 | 10 | 21 | Splicesosom | methylosome, precatalytic spliceosome, SMN-Sm protein complex, U1 snRNP, U5 snRNP |
| 6 | 9 | 308 | Vesicular Transport | clathrin adaptor complex, clathrin adaptor activity, TRAPPIII protein complex, TRAPP complex, Golgi vesicle transport |
| 7 | 8 | 203 | Chemotaxis | C-C chemokine binding, C-C chemokine receptor activity, monocyte chemotaxis, chemokine activity, cell chemotaxis |
| 8 | 8 | 48 | Water/Urea Transport | cellular water homeostasis, glycerol channel activity, urea transmembrane transport, urea transmembrane transporter activity, glycerol transport |
| 9 | 13 | 83 | Signal Transduction | cellular response to cGMP, sphingomyelin catabolic process, sphingomyelin phosphodiesterase activity, Tie signaling pathway, cellular response to macrophage colony-stimulating factor stimulus |
| 10 | 11 | 239 | Cellular Stress Response | aryl hydrocarbon receptor complex, ATF6-mediated unfolded protein response, cellular response to leucine starvation, leucine zipper domain binding, regulation of transcription from RNA polymerase II promoter in response to oxidative stress |
| 11 | 99 | 1337 | Miscellaneous | 1-phosphatidylinositol binding, activation of phospholipase D activity, alpha-2A adrenergic receptor binding, androgen binding, arginine biosynthetic process |
| 12 | 13 | 64 | Sterol Metabolism | 9-cis-retinoic acid biosynthetic process, alcohol dehydrogenase (NADP+) activity, C21-steroid hormone metabolic process, vitamin D catabolic process, cholesterol catabolic process |

To characterize protein families driving these effects, we grouped proteins within the three most prominent clusters (lowest PI-score) based on their UniProt family annotations. Cluster 1 (oxidative degradation) predominantly featured proteins from the cytochrome P450 family. Cluster 4, consistently the most affected, was enriched with potassium channel family proteins, aligning with the GO terms associated with this cluster. Interestingly, cluster 6, linked to vesicular transport, primarily included members of the small GTPase superfamily, such as Rab and Arf family proteins, but also membrane fusion proteins including synaptobrevin and syntaxin family proteins (*Fig. 3E*). This highlights the important role of small GTPases for ATG9A trafficking in AP-4 deficient cells, something that we have also identified in a previous small molecule screen.^33^

Next, we integrated the pathway analysis findings with the results of the secondary screen. Approximately one third of the confirmed hits from the secondary screen were associated with significant pathways, spanning 8 clusters (*Fig. 4A*). Certain genes were exclusively linked to a single pathway cluster, such as *ARL5B* (‘vesicular transport’), *ATF6B* (‘cellular stress response’), *HEATR5B* (‘vesicular transport’), *KCNQ4* (‘potassium channel activity’), *NR3C1* (‘histone H3 demethylation’, *NUTF2* (‘vesicular transport’), *TDP1* (‘signal transduction’) or *TNPO1* (‘vesicular transport’). Others, such as *ADORA1* and *NPM1*, were connected to multiple clusters. Given the cellular phenotypes of AP4-deficiency syndrome, genes involved in vesicular transport (cluster 6) were of particular interest. Notably, *ARL5B*, *HEATR5B*, *NPM1, NUTF2* and *TNPO1* were associated with this cluster. Cluster 2 (‘lipid metabolism’) was represented by *ADORA1*.

**Figure 4:**
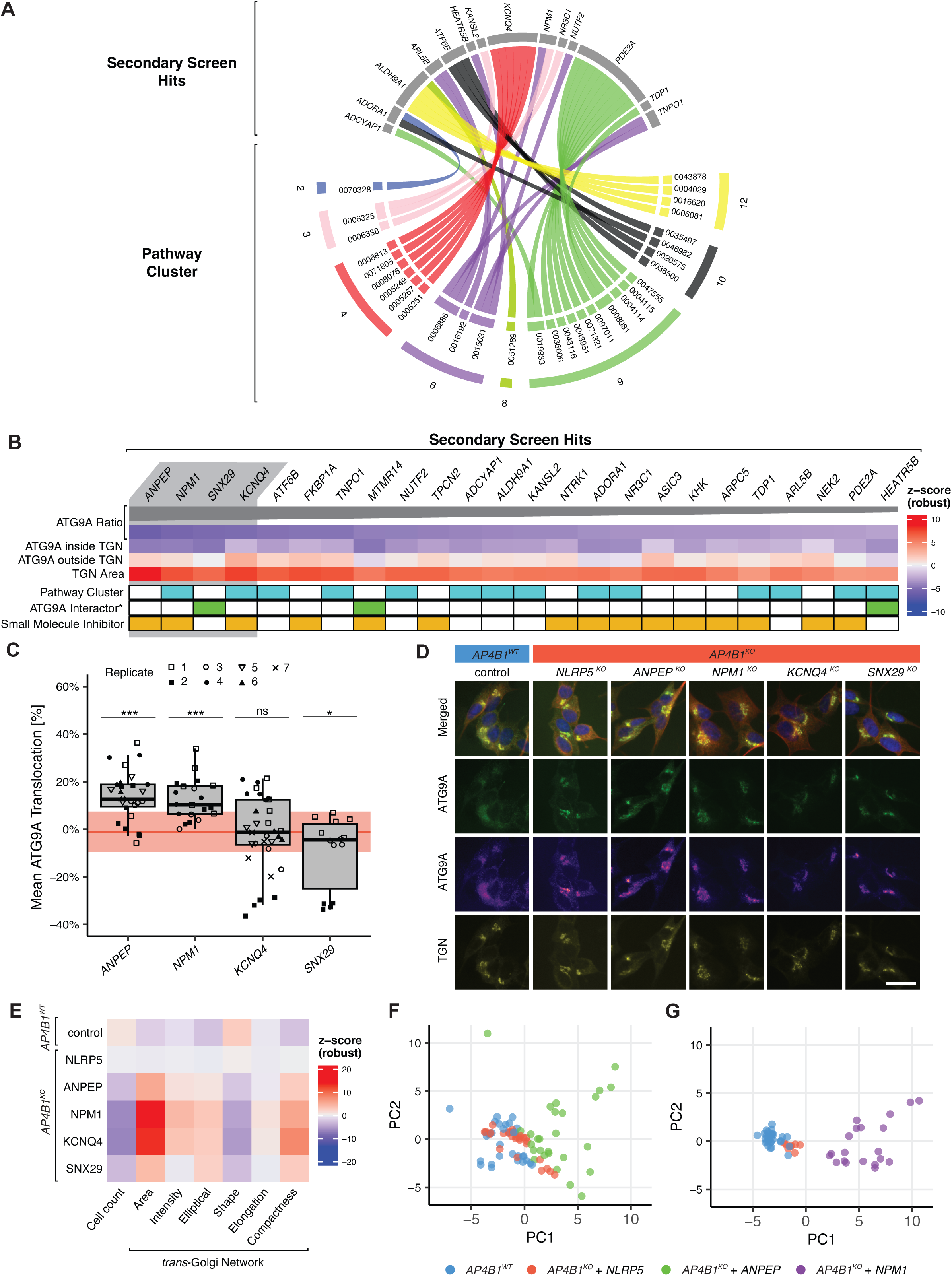
Selection and Validation of Hits Affecting ATG9A Translocation. *A)* Secondary screen hits matched to significantly affected pathways and respective clusters (*Supplementary File 5*), identifying key modulators of ATG9A trafficking in AP4-deficient cells. *B)* Selected secondary screen hits ranked by mean ATG9A ratio reduction across all replicates in the secondary screen and meeting at least one of the following criteria: (1) Association with a pathway cluster, (2) identification as an ATG9A interactor by BioID,^34^ and (3) availability of a specific small molecule inhibitor. Heatmap summarizes secondary screen results. *C)* Mean ATG9A translocation across up to seven independent biological replicates after knockout of the top four genes (*ANPEP*, *NPM1*, *KCNQ4*, and *SNX29*) compared to knockout of the non-essential gene *NLRP5* (red line and area indicate mean ± SD) in AP4B1^KO^ cells. Statistical analysis using one-way ANOVA revealed a significant effect of gene knockout on ATG9A translocation (F(5) = 20.48, p < 0.001, η² = 0.27). Dunnett’s post hoc test indicated significant differences in ATG9A translocation for *AP4B1^KO^* + *ANPEP*, *AP4B1^KO^* + *NPM1*, and *AP4B1^KO^* + *SNX29* compared to *AP4B1^KO^* + *NLRP5*. No significant differences were observed between *AP4B1^KO^* + *KCNQ4* and *AP4B1^KO^* + *NLRP5*. Significance level denoted as p > 0.05 (ns), p < 0.05 (*) and p < 0.001 (***). *D)* Representative fluorescence images of *AP4B1^WT^* and *AP4B1^KO^* cells following knockout of *NLRP5* (negative control), *ANPEP*, *KCNQ4*, and *SNX29*. Images highlight differences in ATG9A localization and TGN morphology. Scale bar = 20 µm. *E)* Heatmap illustrating mean z-scores for cell count and six selected parameters describing TGN morphology. *F)* Dimensionality reduction of 25 ATG9A independent cell morphology parameters by principal component analysis (PCA) demonstrated little to no morphological difference between *AP4B1^WT^* and *AP4B1^KO^* cells. Knockout of *NPM1* resulted in a pronounced deviation from controls, while knockout of *ANPEP* caused substantially less morphological changes.

To corroborate our findings, we compared the hits from the secondary screen with a recent study, which identified ATG9A interactors using quantitative BioID proteomics.^34^ Remarkably, three of our confirmed hits - *HEATR5B*, *MTMR14*, and *SNX29* - were also present in this dataset, highlighting their potential for follow-up studies. Additionally, we evaluated the availability of specific pharmacological inhibitors for the selected genes, facilitating potential clinical translation. We narrowed the list of secondary screen hits to those meeting at least one of the following criteria: (1) association with a relevant pathway cluster, (2) identification as an ATG9A interactor by BioID^34^, and (3) availability of a specific small molecule inhibitor (*Fig. 4B*). The top four hits - *ANPEP*, *NPM1*, *SNX29*, and *KCNQ4* - were prioritized for further validation experiments.

### Validation Experiments Confirm *ANPEP* and *NPM1* as Potential Therapeutic Targets

To ensure the robustness of our findings from the primary and secondary screens, we performed validation experiments with multiple replicates based on nucleofection, a transfection method that achieves higher knockout efficiencies compared to lipofection used in the earlier screens. Knockout efficiency was confirmed via sequencing and analysis with the ICE tool provided by Synthego, which demonstrated knockout scores ranging from 80% to 100%.

Reevaluation of *ANPEP* and *NPM1* knockout confirmed their impact on the ATG9A ratio, consistent with the results from the primary and secondary screens. Conversely, knockout of *KCNQ4* and *SNX29* did not demonstrate a similar ATG9A translocation as seen in both screens (*Fig. 4C, D*). These findings further support the role of ANPEP and NPM1 in regulating ATG9A trafficking in AP4-deficient cells. A detailed analysis of various parameters of TGN integrity revealed less pronounced changes upon knockout of *ANPEP* compared to *NPM1* (*Fig. 4E*), suggesting that *ANPEP* plays a more specific role than *NPM1* in ATG9A trafficking without fundamentally disrupting TGN structure.

To assess the broader effects of *ANPEP* and *NPM1* knockouts on cell morphology, we conducted multiparametric profiling using 25 parameters characterizing the cytoskeleton (i.e. cell shape), nucleus and TGN. Dimensionality reduction by principal component analysis (PCA) across these cellular morphology metrics revealed substantial differences between the two knockouts. *ANPEP* knockout exhibited considerably less variability in cellular morphology compared to *NPM1* knockout, underscoring the lower toxicity associated with *ANPEP* disruption (*Fig. 4F*). This finding highlights *ANPEP* as a promising target for further therapeutic exploration, with a more favorable toxicity profile than *NPM1*.

## DISCUSSION

Despite considerable progress in recent years, the molecular mechanisms underlying AP-4-associated HSP remain incompletely understood. Deciphering these cellular processes is critical for developing new therapeutic strategies for this progressive childhood-onset disease. In this study, we conducted a phenotypic high-content arrayed CRISPR/Cas9-based loss-of-function screen in a neuronal model of AP-4 deficiency, revealing key pathways and identifying modulators of ATG9A trafficking in the context of AP-4 deficiency.

CRISPR/Cas-based functional genomics screens have proven to be a powerful tool for elucidating gene and pathway function. Leveraging the advantages of CRISPR technology over previous genomic screening methods, we interrogated a library targeting approximately one-third of the human genome. To achieve this, we developed an arrayed screening platform using a neuronal cell model of AP-4 deficiency syndrome. By combining this platform with imaging-based multiparametric single-cell analysis, we demonstrated the strengths of arrayed CRISPR screening - offering high accuracy in gene function interrogation alongside a custom experimental and analytical pipeline. A robust assay design, including multiple replicates, stringent controls, and extensive automation, allowed us to maximize the utility of this approach.^35^

However, we acknowledge certain limitations in our study. For instance, the alphabetical arrangement of the sgRNA library in the primary screen introduced potential plate-based artifacts. Nonetheless, by implementing strict quality criteria and conducting a secondary screen with randomized plate positions, we minimized these risks. Another limitation is the transfection efficiency achieved using lipofection, which may have led to false negatives for genes with small effect sizes. Despite these constraints, our approach has successfully uncovered several regulators of ATG9A distribution in AP-4 deficient cells.

Our pathway analysis identified several clusters and protein classes as critical modulators of AP4-deficiency syndrome, offering potential therapeutic targets. As anticipated, proteins involved in vesicular transport emerged as key regulators. Specifically, we identified small GTPases from the Rab and Arf families as essential regulators.

This aligns with previous studies implicating small GTPases in ATG9A and AP-4 cycling. For example, Rab7 has been shown to regulate the recruitment of ATG9A-positive vesicles for mitophagy,^36^ while Rab1 is essential for autophagosome formation and is present on ATG9A vesicles.^37^ Beyond this, Rab10 is necessary for ATG-9 trafficking in *Caenorhabditis elegans*.^38^ Our group previously demonstrated that small molecules targeting Rab3C and Rab12 can restore ATG9A distribution in AP-4-deficient cells.^33^ Arf proteins also play crucial roles; for instance, Arf1 mediates AP-4 localization to the TGN in yeast.^39^ However, recent studies suggested that in mammalian cells, this function may be carried out by the Arf-like protein Arl5b, which also regulates retrograde endosome-to-TGN transport.^40,41^ In our screen, loss of Arl5b significantly promoted ATG9A redistribution. Potentially, this is mediated by diminished retrograde transport of ATG9A to the TGN.

Another hit, *NR3C1*, encodes for a nuclear glucocorticoid receptor and is related to the significant pathway cluster ‘histone H3 demethylation’. In mouse pancreatic β cells, NR3C1 was found to enhance autophagy via increased expression of the RNA demethylase FTO (fat mass and obesity-associated protein). FTO catalyzes m^6^A demethylation of *Atg9a* which promotes mRNA stability and thereby upregulates *Atg9a* expression.^42^ Consequently, knockout of *NR3C1* in our screen likely lowered *ATG9A* expression and reduced its accumulation in the TGN.

The conservation of AP-4-mediated vesicular transport across eukaryotes is striking; plants exhibit similar cellular phenotypes to mammalian cells in the context of AP-4 deficiency.^43,44^ In *Arabidopsis thaliana* seedlings, AP-4 loss depleted cytochrome P450 proteins.^44^ In our pathway analysis, cluster 1 dominated by cytochrome P450 enzymes was among the top three clusters, suggesting a conserved role for these enzymes in AP-4-dependent vesicular trafficking. Additionally, lipid metabolism pathways (cluster 2) emerged as relevant. The diverse interconnections between AP-4 and lipid metabolism have been established in both human cells and mouse models, i.e. by the identification of AP-4 cargos such as low-density lipoprotein receptor (LDLR) and diacylglycerol lipase beta (DAGLB).^16,45^ Furthermore, altered trafficking of the AP-4 cargo ApoER2 (apolipoprotein E receptor 2) in AP-4 deficient cells implicated aberrant Reelin signaling.^46^

Interestingly, our screen uncovered cation channels, particularly potassium channels, as modulators of ATG9A cycling. Two hits from the secondary screen, *KCNQ4* and *TPCN2*, are prominent representatives. TPC2, a non-selective cation channel localized to endolysosomal membranes encoded by the *TPCN2* gene, is involved in intracellular transport and lysosomal exocytosis, and has been investigated as a therapeutic target for lysosomal storage diseases and neurodegenerative disorders.^47^ However, KCNQ4, a potassium channel critical for neuronal excitability in cochlear sensory cells,^48^ together with SNX29, a cellular trafficking protein with a phospholipid-binding domain^49^ that has been suggested to be associated with mental disorders^50^, showed inconsistent results upon repeated testing. This highlights the challenges of validating subtle phenotypic changes in high-content screens.

Among the confirmed hits, *ANPEP* (alanyl aminopeptidase) and *NPM1* (nucleophosmin) emerged as strong candidates for further exploration. Knockout of both genes consistently influenced ATG9A redistribution. However, the impact of *ANPEP* knockout on TGN morphology was less pronounced. This suggested a more specific modulation of ATG9A trafficking, whereas *NPM1* knockout implicated a less targeted effect which resulted in TGN disruption, possibly causing unselective cargo escape. The pleiotropic functions of ANPEP are well-documented,^51–53^ but its involvement in TGN protein export represents a novel finding. Combined with its robust effect on ATG9A trafficking and low toxicity, ANPEP represents a potential target for therapeutic investigation.

In summary, our study provides insights into the mechanisms regulating ATG9A trafficking in AP4-deficient neuronal cells. We identified candidate genes, such as *ANPEP*, that warrant further investigation as potential drug targets for AP4-HSP. Additionally, we present a comprehensive dataset for the research community and outline an experimental and analytical pipeline that serves as a blueprint for future arrayed cell-based phenotypic screens.

## RESOURCE AVAILABILITY

### Lead contact

Further information and requests for resources and reagents should be directed to and will be fulfilled by the lead contact, Darius Ebrahimi-Fakhari, MD, PhD.

### Materials availability

This study did not generate new unique reagents.

### Data and code availability

- Screening data have been deposited at Zenodo and are publicly available as of the date of publication. DOIs are listed in the key resources table.
- Relevant original code has been deposited at Zenodo and is publicly available as of the date of publication. DOIs are listed in the key resources table.
- Any additional information required to reanalyze the data reported in this paper is available from the lead contact upon request.

## ACKNOWLEDGEMENTS

The authors thank Jennifer Smith, PhD, Clarence Yapp, PhD, and the team from the ICCB-Longwood Screening Facility for help with designing and conducting screening experiments. This study was supported by research grants from the CureAP4 Foundation (to D.E.-F.), the Spastic Paraplegia Foundation (to D.E.-F.), the Manton Center for Orphan Disease Research (to D.E.-F.), the BCH Office of Faculty Development (to D.E.-F.), the BCH Translational Research Program (to D.E.-F.) and the National Institute of Neurological Disorders and Stroke (1K08NS123552-01 to D.E.-F.). Further support is acknowledged from the German National Academic Foundation (to M.Z., C.B., J.E.A), the Carl Duisberg Fellowship for Medical Sciences by the Bayer Foundation (to M.Z.), the Biomedical Education Program by the German National Exchange Service (to M.Z., C.B., J.E.A), the RWTH-Aachen-Research-Ambassador-Scholarship (to C.B), the German Research Foundation (to A.S. - 448402208), the European Union’s Horizon 2020 research and innovation program under the Marie Sklodowska-Curie grant agreement no. 896725 (to A.K.D), and the Rosamund Stone Zander chair (to M.S.). The IDDRC at Boston Children’s Hospital is supported by National Institutes of Health grant 1U54HD090255.

## AUTHOR CONTRIBUTIONS

M.Z., C.B., J.E.A, A.S., A.K.D., D.E.F. conceptualized and designed the experiments. M.Z., C.B., J.E.A, H.M.K., D.E.F. performed experiments. M.Z. and D.E.F. wrote the first draft of the manuscript. All authors contributed to the final draft of the manuscript. D.E.F. wrote the grants that designed and supported this project. M.S. and D.E.F. supervised the project.

## SUPPLEMENTAL INFORMATION

### Supplementary Figures

Supplementary File 1: Complete dataset for the primary screen

Supplementary File 2: Complete dataset for the secondary screen

Supplementary File 3: List of GO terms derived from Ensemble (accessed: 09/19/2022)

Supplementary File 4: Complete dataset of significant pathways

Supplementary File 5: Significant pathways in relevant pathway clusters associated with secondary screen hits

Supplementary File 6: Raw data for Figures 4 *C*, *E, F* and *G*

## METHODS

### Key resources table

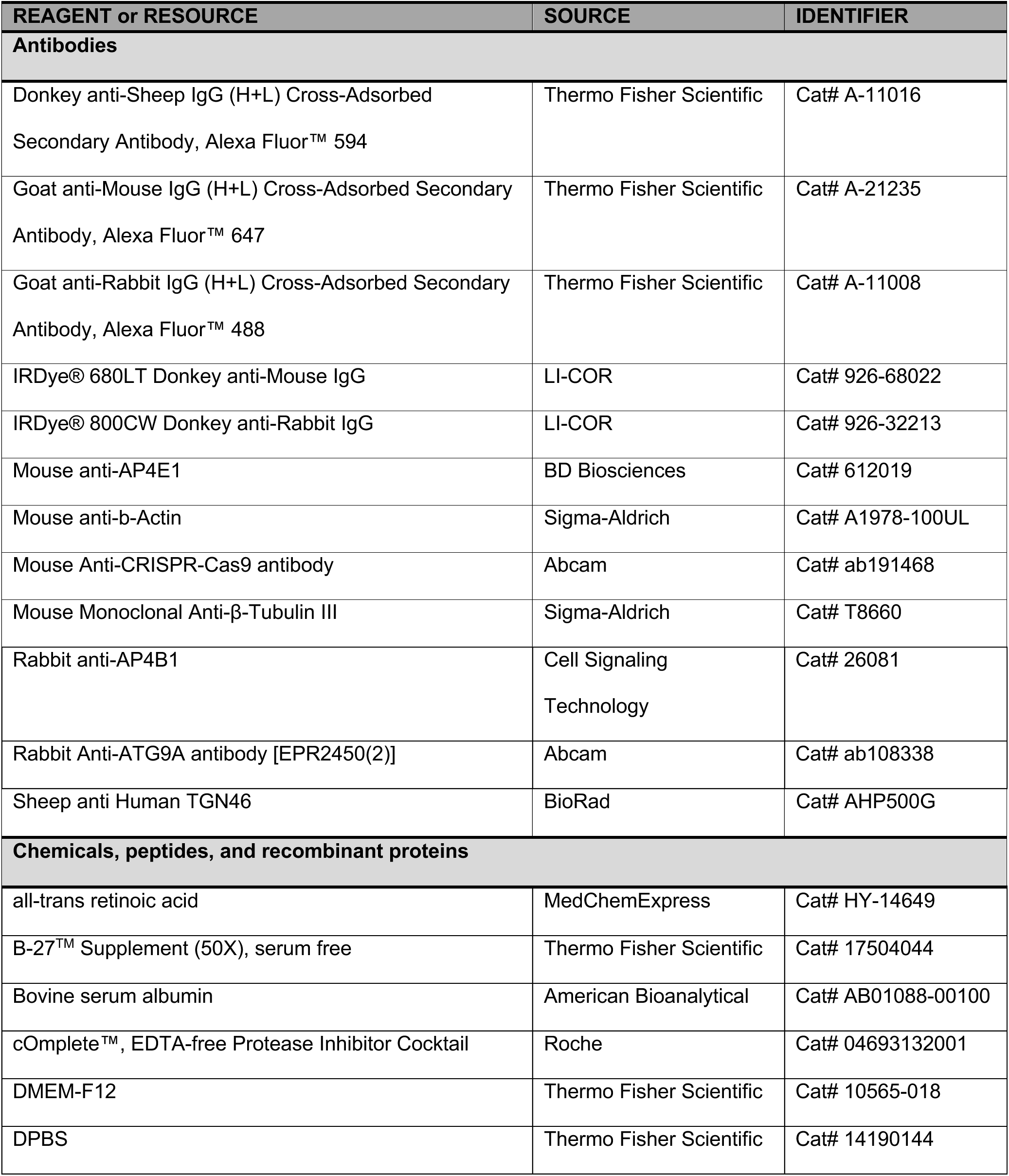
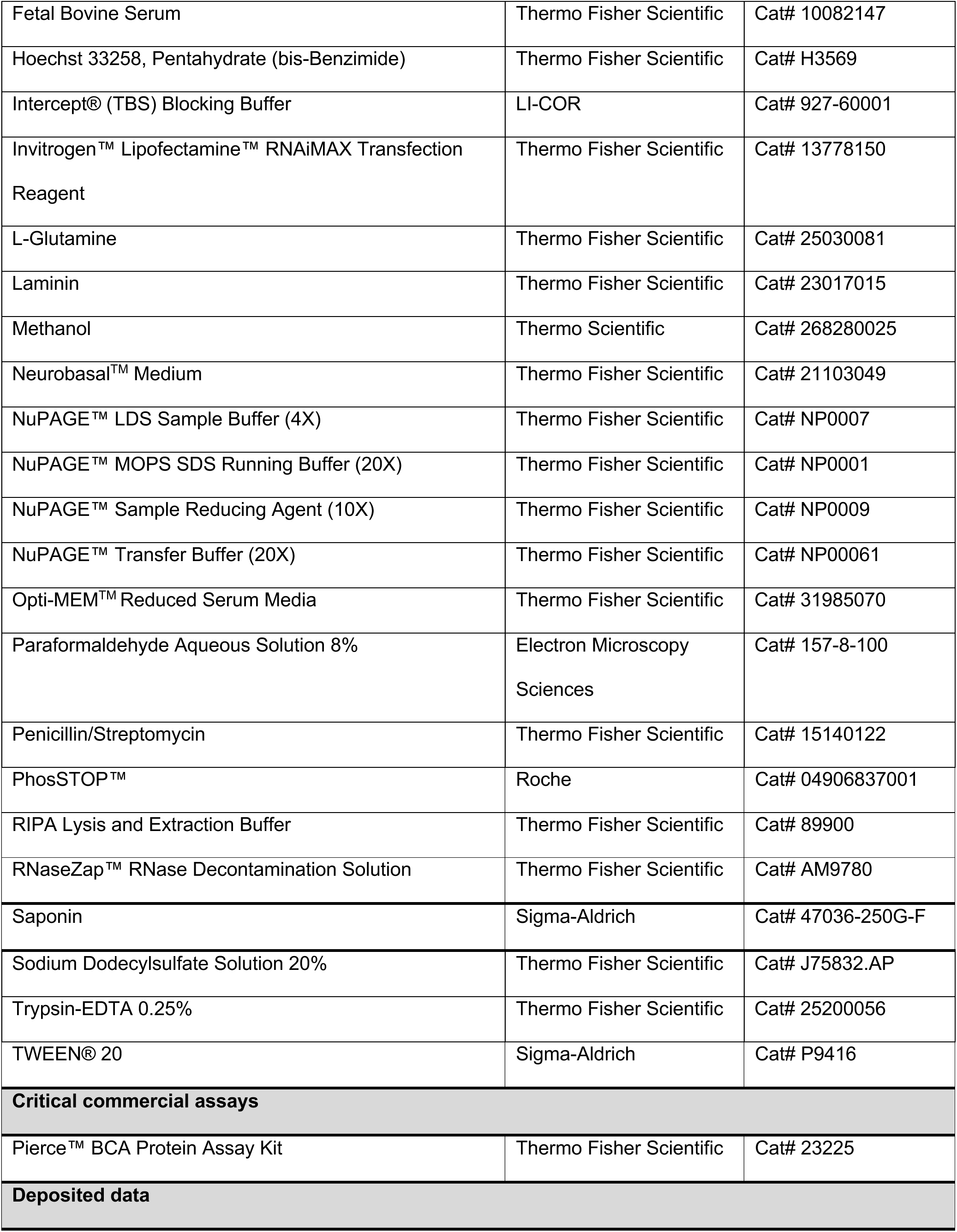
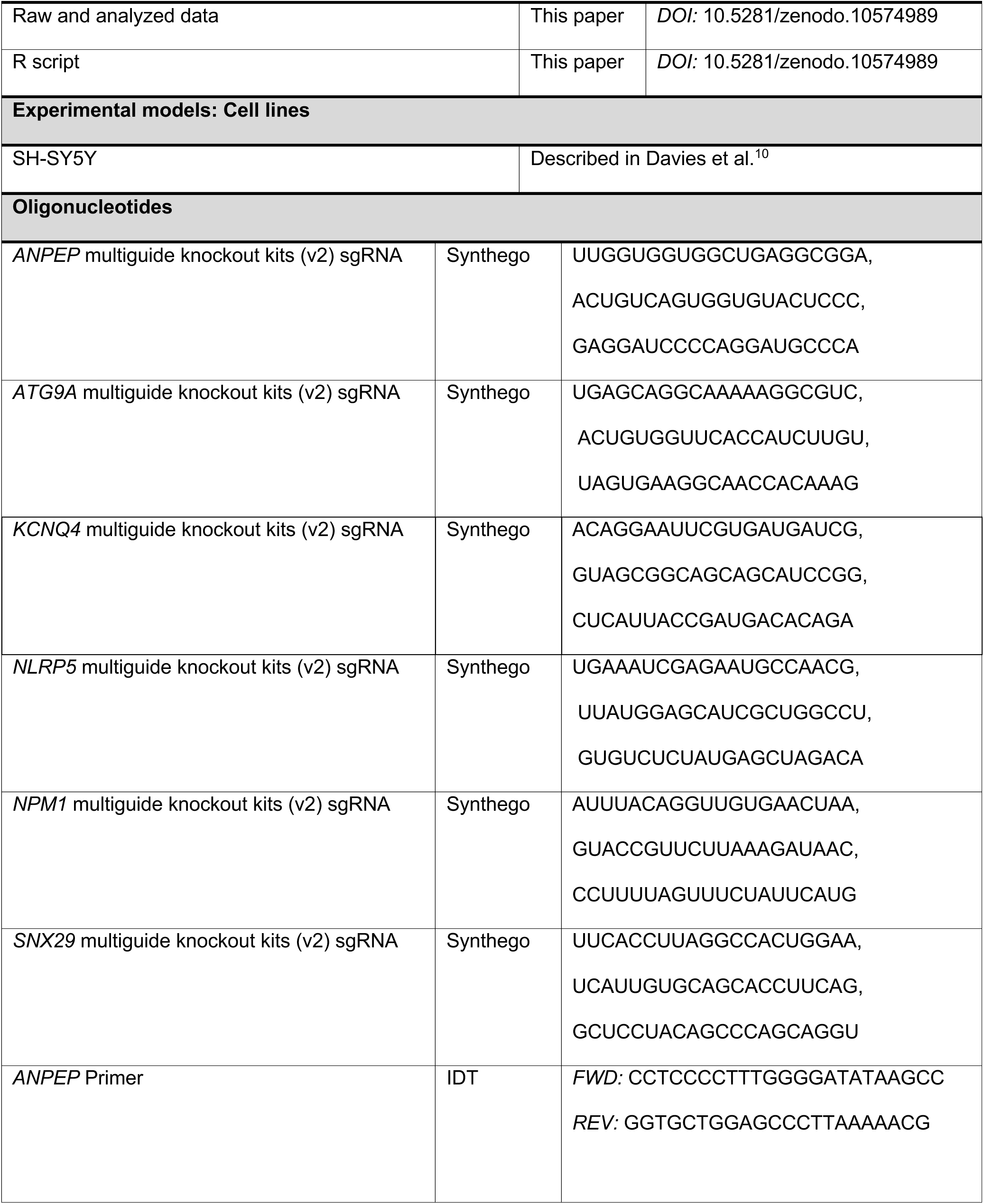
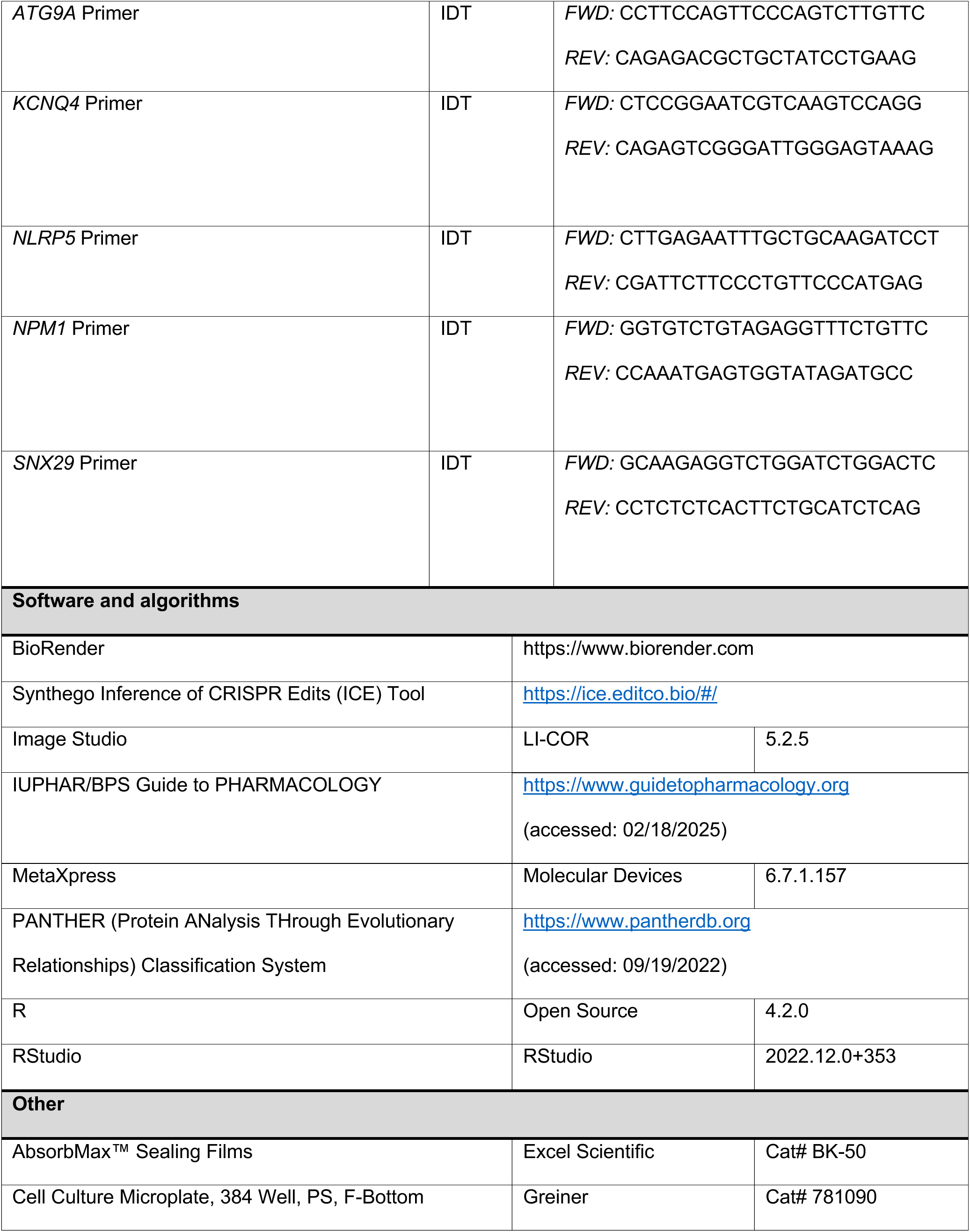
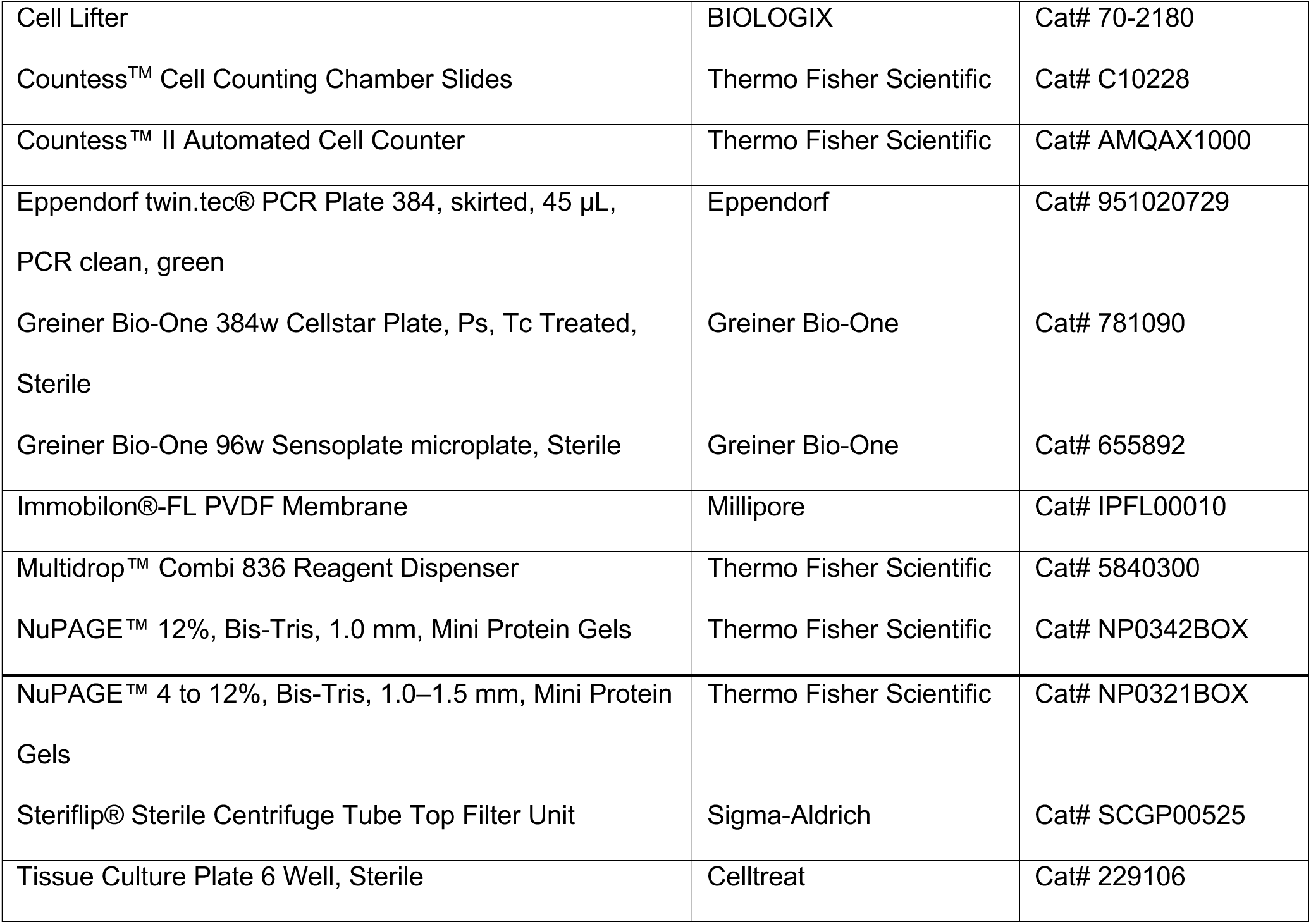

### Experimental model and study participant details

#### SH-SY5Y Cells

For this study, we used previously published SH-SY5Y cells stably expressing a sgRNA targeting *AP4B1* and Cas9, and SH-SY5Y cells stably expressing Cas9 but no sgRNA.^10^ Cells were cultured in Dulbecco’s Modified Eagle Medium/Nutrient Mixture F-12 supplemented with 10% Fetal Bovine Serum (FBS), 100 U/mL penicillin, and 100 µg/mL streptomycin. Cultures were maintained at 37°C in a humidified incubator with 5% CO₂. Cells were passaged every three days by washing with Dulbecco’s phosphate-buffered saline (DPBS) and dissociating with 0.25% Trypsin-EDTA. Media changes were performed every other day to ensure optimal growth conditions. For differentiation, SH-SY5Y cells were cultured in neurobasal medium supplemented with 1× B-27 supplement, 2 mM L-glutamine, and 10 µM all-*trans* retinoic acid for five days. To improve differentiation and enhance cell attachment, laminin (10 µg/mL) was added to the differentiation media.^54^

#### SgRNA Library

The ‘Synthego sgRNA Human Druggable Genome’ library was obtained from the ICCB Longwood Screening Facility at Harvard Medical School. This library targets 8,478 human genes, utilizing three distinct chemically synthesized sgRNAs per gene, each designed to target early exons. The library is arrayed across 29 plates in a 384-well format, with designated wells allocated for assay-specific controls to ensure experimental consistency.

### Method details

#### Reverse Transfection

All steps were conducted in an RNase- and DNase-free environment to ensure sample integrity. Positive and negative control sgRNAs (targeting *ATG9A* and *NLRP5*, respectively) were diluted in nuclease-free Tris-EDTA buffer and dispensed into the appropriate wells of a 384-well PCR plate to achieve a final concentration of 50 nM. Lipofectamine™ RNAiMAX transfection reagent was diluted in reduced-serum medium (Opti-MEM) and distributed into three 384-well assay plates per library plate to create three replicates. The final volume of RNAiMAX in the reaction was adjusted to 0.16%. The sgRNA library and control sgRNAs were transferred to the RNAiMAX-containing assay plates using the Agilent Bravo Automated Pipettor and the Agilent 4-R BenchCel Plate Handler. SH-SY5Y cells cultured in differentiation media for two days were counted using an automated cell counter. Cells were then plated at a density of 3,000 cells per well in differentiation media containing 10 µM all-*trans* retinoic acid, but without laminin. The Multidrop™ Combi Reagent Dispenser was employed for RNAiMAX solution dispensing and cell plating. Plates were spun briefly to ensure even distribution of reagents and incubated at room temperature for 20 minutes to allow uniform cell settlement. The assay plates were incubated for 72 hours at 37°C in a 5% CO₂ humidity-controlled incubator to minimize evaporation and edge effects. After 24 hours of incubation, 10% differentiation media supplemented with 40 µM all-*trans* retinoic acid and 10 µg/mL laminin was added to support differentiation and cell adhesion.

#### Nucleofection

All steps were conducted in an RNase- and DNase-free environment to maintain sample integrity. To enhance gene-editing efficiency, ribonucleoprotein complexes (RNPs) were utilized for nucleofection. sgRNAs were diluted in TE buffer to a stock concentration of 100 µM, and Cas9 was prepared at a stock concentration of 20 µM. The nucleofector solution was prepared by mixing the nucleofector reagent and supplement at a 4.5:1 ratio. RNPs were assembled by combining sgRNA (final concentration: 7.2 µM) and Cas9 protein (final concentration: 0.8 µM) in the nucleofector solution. Briefly, 180 pmol sgRNA was incubated with 20 pmol Cas9 protein in the nucleofector solution for at least 10 minutes to form RNP complexes. SH-SY5Y cells were harvested, and 4 × 10⁵ cells were resuspended in 5 µL of the prepared nucleofector solution. The cell suspension was mixed with the pre-assembled RNPs at a 1:6 ratio (cell suspension/RNP solution) and transferred into nucleofection strips. The strips were placed in the 4D-Nucleofector System, and nucleofection was carried out using the CA-137 program. After nucleofection, pre-warmed medium was added to the cells following a 10-minute incubation at room temperature. The cells were subsequently plated at a density of 1 × 10⁴ cells per well into a 96-well plate using the Integra Voyager 8-channel automated pipettor.

#### Fixation and Immunocytochemistry

After 72 hours of incubation, cells were fixed with 4% paraformaldehyde for 20 minutes at room temperature. Cells were then permeabilized using 0.1% saponin in DPBS for 10 minutes. Following permeabilization, the solution was removed and blocking buffer containing 0.1% bovine serum albumin (BSA) and 0.01% saponin in DPBS was added for 20 minutes to minimize non-specific binding. Primary antibodies were diluted in blocking buffer to the following concentrations: rabbit anti-ATG9A (1:500 to 1:1000), sheep anti-TGN46 (1:800), and mouse anti-β-Tubulin III (1:1000). Cells were incubated with primary antibodies for 1 hour at room temperature and then washed three times with blocking buffer. Secondary antibodies were prepared in blocking buffer at concentrations of 1:1250 to 1:2000 and included Alexa Fluor 488-conjugated anti-rabbit, Alexa Fluor 594-conjugated anti-sheep, and Alexa Fluor 647-conjugated anti-mouse. Additionally, Hoechst 33258 (1:2000) was used for nuclear staining. Cells were incubated with secondary antibodies for 30 minutes in the dark, followed by three washes with DPBS to remove unbound antibodies. All solutions used for immunostaining were filtered through a 0.22 µM filter system to ensure sterility and consistency. After staining, plates were sealed with opaque seals to prevent light exposure and prepared for imaging.

#### High-Throughput Imaging

High-throughput imaging in 384-well plates was performed using the Molecular Devices ImageXpress Micro Confocal Laser System, equipped with a 40× S Plan Fluor objective (numerical aperture: 0.60 µm, working distance: 3.6–2.8 mm). For each well, 16 fields were acquired in a 4 × 4 grid configuration. For high-throughput imaging in 96-well plates, 24 sites per well were captured in a 5 × 5 grid configuration, leaving out the center position. Sites were spaced 330 µm apart to ensure even coverage of each well. A PAA GX™ robot arm was used for automated plate loading into the microscope. Image analysis was conducted using a customized pipeline within the MetaXpress software (Molecular Devices, version 6.7.1.157). Cells were identified as β-Tubulin III-positive areas containing Hoechst-positive nuclei. The *trans*-Golgi network (TGN) was defined by TGN46 signal intensity, while ATG9A protein was identified based on its specific signal intensity. Overlapping TGN46 and ATG9A signals were classified as ‘ATG9A inside the TGN’, whereas ATG9A signal outside the TGN46-defined region but within the cellular boundary was categorized as ‘ATG9A outside the TGN’.

#### Western Blotting

Cells were cultured in 6-well plates at a density of 5 × 10⁵ cells per well and harvested using RIPA buffer supplemented with protease inhibitors and phosphatase inhibitors. The total protein concentration was determined using the Pierce™ BCA Protein Assay Kit with absorbance measured at 562 nm using a Tecan plate reader. Protein samples were adjusted to a total amount of 15 µg, mixed with NuPAGE™ LDS Sample Buffer, and supplemented with NuPAGE™ Sample Reducing Agent. The samples were boiled at 70°C for 10 minutes before loading onto NuPAGE™ 4–12% Bis-Tris gels. Electrophoresis was carried out in NuPAGE™ MOPS SDS Running Buffer for approximately 50 minutes at 200 V. Proteins were transferred onto an Immobilon®-FL PVDF Membrane using a wet transfer system with NuPAGE™ Transfer Buffer at 30 V for 75 minutes. The membrane was then blocked with Intercept® (TBS) Blocking Buffer for 60 minutes on a shaker. Membranes were incubated overnight at 4°C with the following primary antibodies, diluted in Intercept® (TBS) Blocking Buffer supplemented with 0.2% TWEEN® 20:

- Mouse anti-CRISPR-Cas9 (1:500) (two-day incubation),
- Rabbit anti-AP4B1 (1:500),
- Mouse anti-β-Actin (1:2000),
- Mouse anti-AP4E1 (1:500),
- Rabbit anti-ATG9A (1:500).

Near-infrared secondary antibodies (IRDye® 800CW Donkey anti-Rabbit IgG; IRDye® 680LT Donkey anti-Mouse IgG) were diluted 1:5000 in Intercept® (TBS) Blocking Buffer supplemented with 0.2% TWEEN® 20 and 0.01% sodium dodecyl sulfate. Membranes were incubated with secondary antibodies for 1 hour at room temperature in the dark. Protein detection and imaging were performed using the LI-COR Odyssey® DLx system. Image analysis was carried out using the Image Studio software (LI-COR, version 5.2.5).

#### Available Small Molecule Inhibitors

Available small molecule inhibitors for prioritizing secondary screen hits (*Fig. 4B*) were identified using the ‘International Union of Basic and Clinical Pharmacology (IUPHAR) / British Pharmacological Society (BPS) Guide to Pharmacology’ (https://www.guidetopharmacology.org, accessed: 02/18/2025) and the product portfolio of MedChemExpress® (https://www.medchemexpress.com, accessed: 02/18/2025).

### Quantification and statistical analysis

Statistical analysis and plotting were performed using R statistical computing (version 4.2.0) and the RStudio IDE (version 2022.12.0+353). The following R packages were used: dplyr,^55^ ggplot2,^56^ openxlsx,^57^ ggpubr,^58^ plotly,^59^,ggrepel,^60^ ggExtra,^61^ janitor,^62^ lsr,^63^ tidyr,^64^ tibble,^65^ ComplexHeatmap,^66,67^ multcomp,^68^ effectsize,^69^ bibtex,^70^ UniprotR,^71^ dynamicTreeCut,^72^ ggbreak,^73^ gggap,^74^ pbapply,^75^ cluster,^76^ purrr,^77^ moduleColor,^78^ pals,^79^ Hmisc,^80^ biomaRt,^81,82^ scales,^83^ RColorBrewer,^84^ circlize.^85^

For normalization, the z-score robust was calculated:

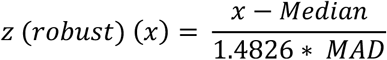

(x: value per well, Median: median of all wells, MAD: median absolute deviation, 1.4826: constant to adjust MAD for asymptomatically normal consistency)

As a quality metric for assay plates to account for plating artefacts and staining irregularities, we calculated the Z’ factor and the SSMD as well as the robust version of both parameters using the median and median absolute deviation.

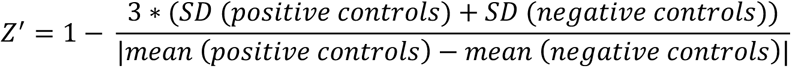

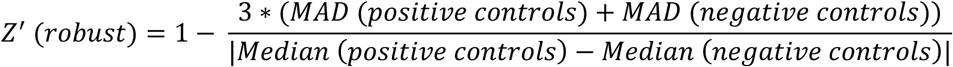

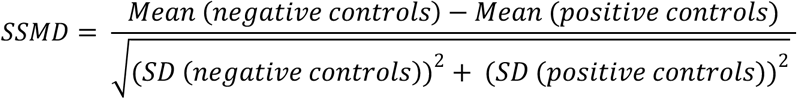

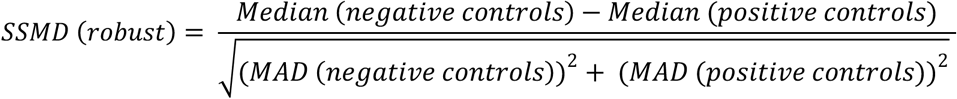

Data is summarized by mean ± standard deviation. The Pearson correlation coefficient was calculated between all replicates using the *cor()* function in R. Statistical differences between two groups were evaluated using the Student’s t-test, with p-values < 0.05 considered statistically significant. Effect sizes for two-group comparisons were estimated using Cohen’s d, with the following thresholds applied: a Cohen’s d of ≤0.2 indicated a small effect, 0.5 a medium effect, and ≥0.8 a large effect.^86^ For comparisons involving multiple groups, one-way analysis of variance (ANOVA) was used, followed by Tukey’s HSD or Dunnett’s multiple comparison post-hoc test for comparisons to the control group. Effect sizes for multiple group comparisons were estimated using eta squared (η²), with thresholds defined as: η² ≤ 0.01 indicating a small effect, η² = 0.06 a medium effect, and η² ≥ 0.14 a large effect.^86^ A total of three wells in the *AP4B1^KO^* + *NLRP5* control group were excluded from analysis due to screening artifacts.

The morphological profile of the TGN was analyzed using TGN roughness (shape factor in the MetaXpress software) and the following calculated metrics:

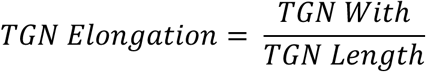

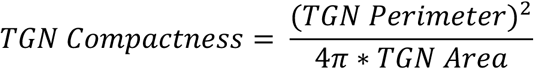

For pathway analysis, a list of GO terms associated with genes from the screen was derived from Ensembl (*Supplementary File 3*). GO terms with less than 3 or more than 1,000 genes in our screen were excluded. For each GO term, a Pathway Impact Score (PI Score) was calculated.^30^

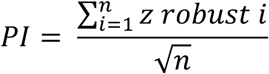

(PI: Pathway Impact Score, n = number of genes associated with GO term, z robust i: z-score robust for each gene associated with GO term)

The Pathway Impact Score calculation for each GO term size, i.e. the number of genes comprised by each GO term, was performed 10,000 times using randomly selected values from the screen for the z-score robust. From this, the normal distribution of PI scores for each GO term could be derived and the probability p of obtaining the actual PI score by chance was calculated. Similar sized GO terms yielded a similar normal distribution. Therefore, p-values were adjusted for multiple testing according to Bonferroni by multiplication with the number of occurrences of the respective GO term size. A pathway was considered significantly affected if p adjusted < 0.05.

To determine the overlap between two pathways, we calculated the overlap coefficient.^30^

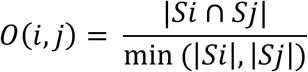

|*Si ∩ Sj*| is the number of shared genes between the two gene sets Si and Sj, min (|*Si|,|Sj*) is the size of the smaller set.

A matrix of overlap coefficients was calculated for all pathways. From this, a distance matrix based on Euclidean distance was generated, followed by average linkage clustering. The hierarchical clustering order was applied to reorder the overlap coefficient matrix, and a heatmap view was created for visualization. Clusters of highly overlapping pathways were identified using the ‘Dynamic Tree Cut’ algorithm.^72^ Suitable umbrella terms were manually assigned to each cluster to provide descriptive labels. For the proteins associated with each cluster, family domain information was retrieved from the Universal Protein Resource (UniProt).^71^

## Additional resources

The Synthego sgRNA Human Druggable Genome Library is available through the ICCB-Longwood Screening Facility at Harvard Medical School: https://iccb.med.harvard.edu/synthego-sgrna-human-druggable-genome

Detailed nucleofection protocol for Lonza 4D Nucleofector: https://bioscience.lonza.com/lonza_bs/DE/en/document/21324

