## Supplemental Files for "Arrayed CRISPR/Cas9 Loss-Of-Function Screen in a Neuronal Model of Adaptor Protein Complex 4 Deficiency Identifies Modulators of ATG9A Trafficking"

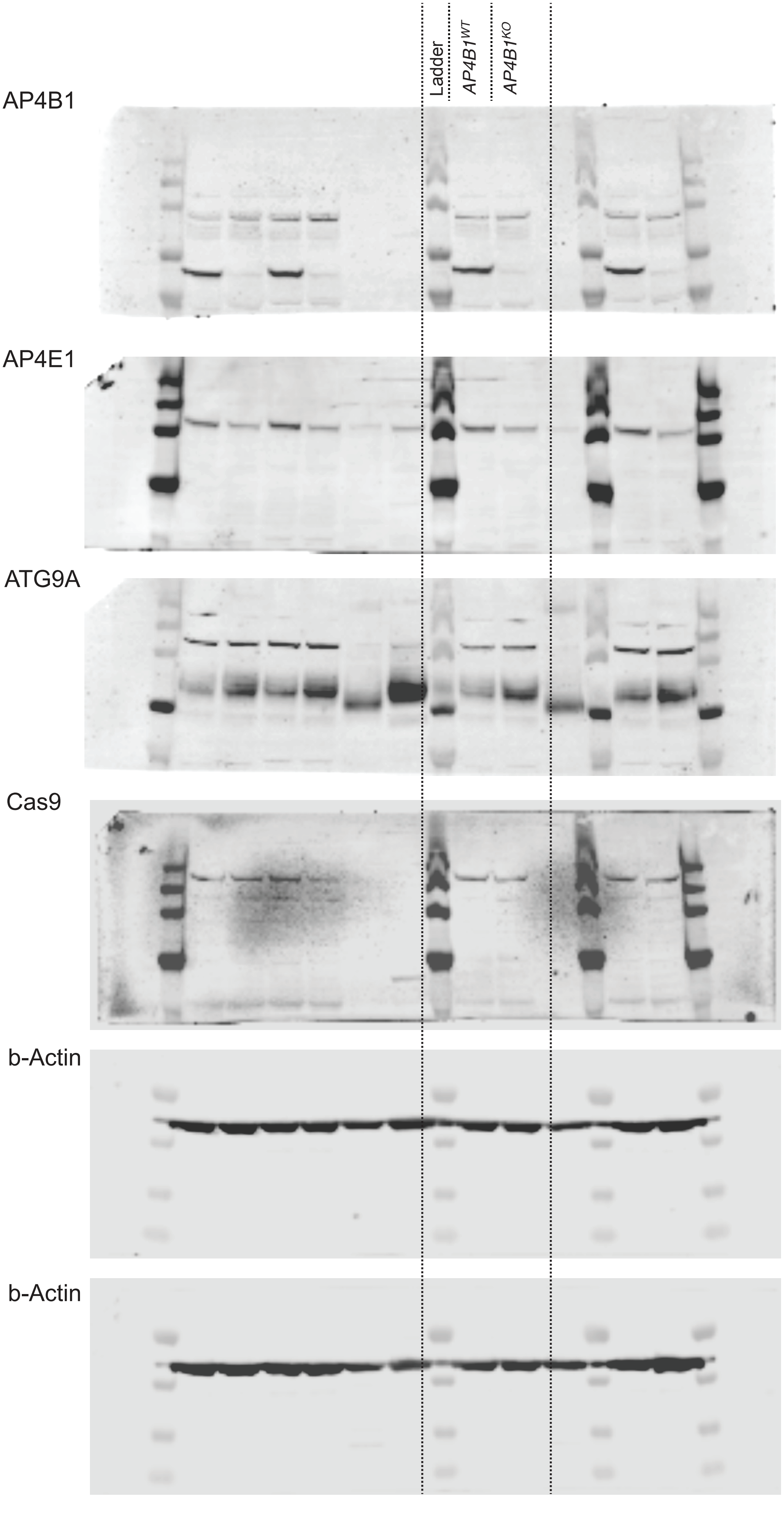
Ladder
AP4B1$^{WT}$
AP4B1$^{KO}$
AP4B1
AP4E1
ATG9A
Cas9
b-Actin
b-Actin

#### Supplementary Figure 1: Uncut Western Blots (*Figure 1B)*

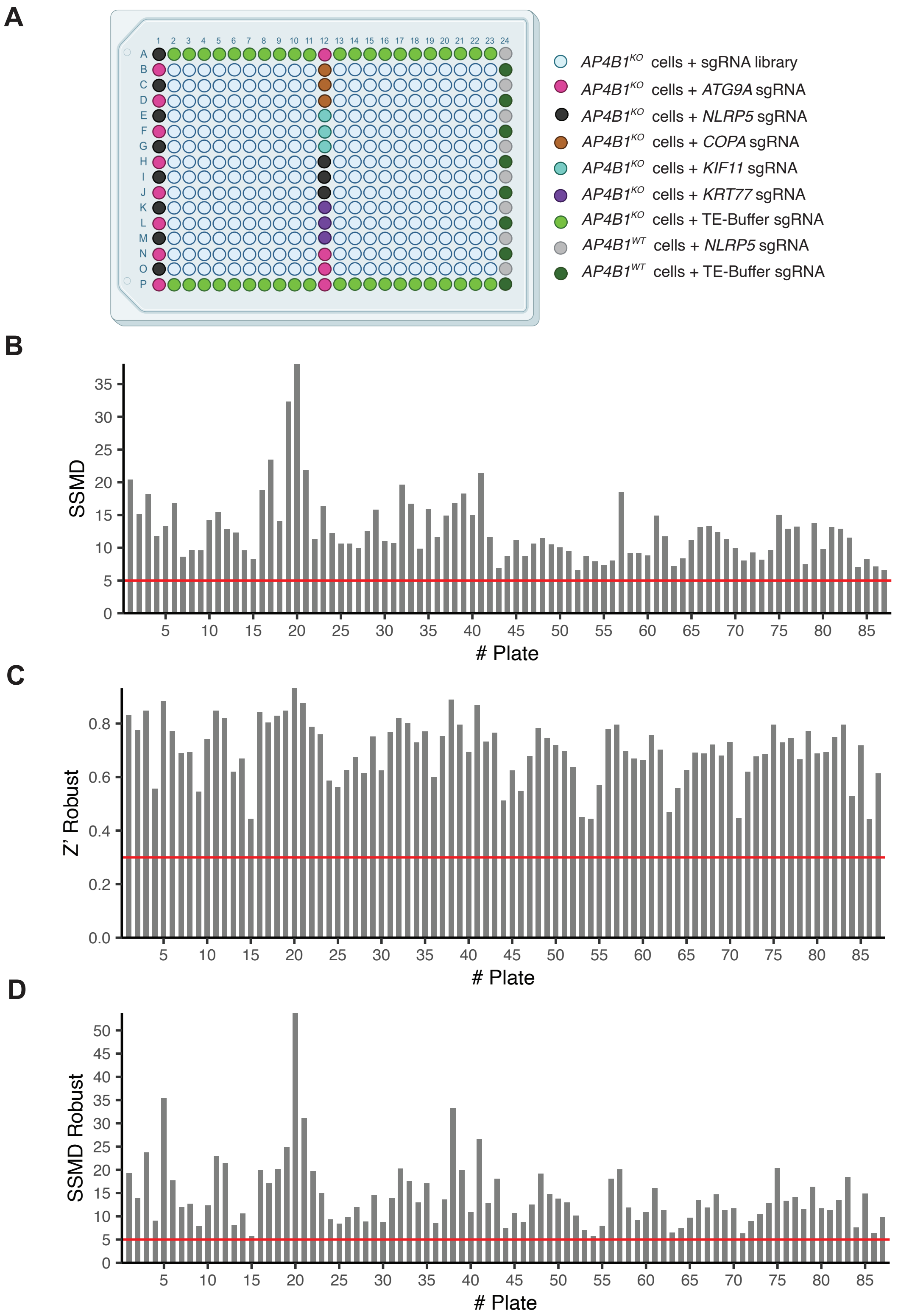
A
AP4B1KO cells + sgRNA library
AP4B1KO cells + ATG9A sgRNA
AP4B1KO cells + NLRP5 sgRNA
AP4B1KO cells + COPA sgRNA
AP4B1KO cells + KIF11 sgRNA
AP4B1KO cells + KRT77 sgRNA
AP4B1KO cells + TE-Buffer sgRNA
AP4B1WT cells + NLRP5 sgRNA
AP4B1WT cells + TE-Buffer sgRNA
B
SSMD
### Plate
C
Z' Robust
### Plate
D
SSMD Robust
### Plate

**Supplementary Figure 2: Primary Screen Plate Design and Quality Control Metrics**

*A)* Plate layout for primary screen. Positive controls included *AP4B1*$^{KO}$ cells transfected with sgRNAs targeting *ATG9A*, and *AP4B1*$^{WT}$ cells treated with TE-buffer or transfected with sgRNAs targeting the non-essential gene *NLRP5*. Negative controls comprised *AP4B1*$^{KO}$ cells treated with TE-buffer or transfected with sgRNAs targeting non-essential genes *NLRP5* or *KRT77*. To assess transfection and knockout efficiency, cell death-inducing sgRNAs targeting essential genes *COPA* and *KIF11* were used. This figure was created with BioRender.com.

*B)* Strictly standardized mean difference (SSMD) for all 87 screened plates demonstrated robust assay performance. Horizontal line indicates predefined quality control threshold (SSMD > 5).

*C)* Z' robust for all 87 screened plates. Horizontal line indicates predefined quality control threshold (Z' > 0.3).

*D)* SSMD robust for all 87 screened plates. Horizontal line represents predefined quality control threshold (SSMD robust > 5).

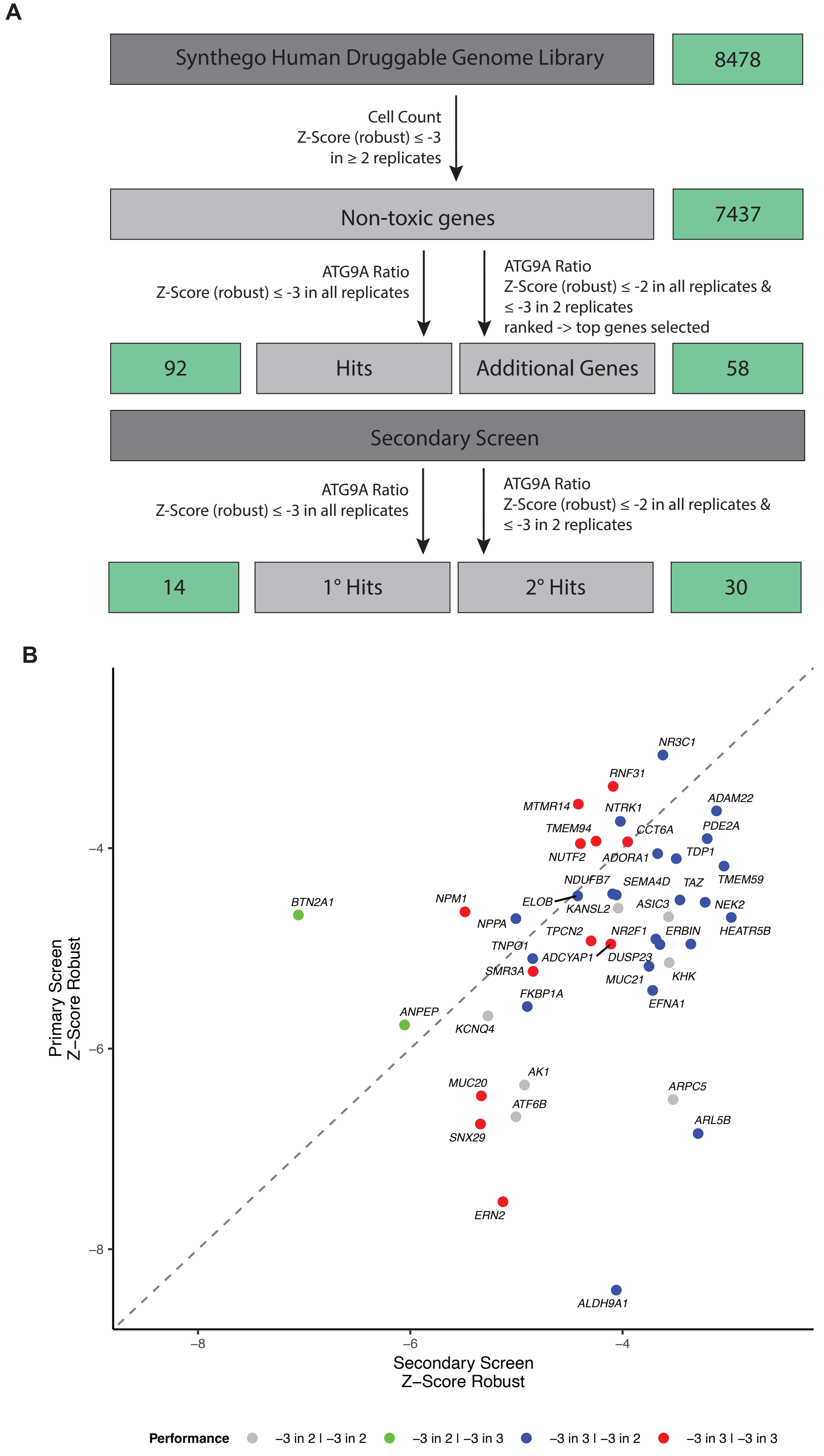
A
Synthego Human Druggable Genome Library
8478
Cell Count
Z-Score (robust) ≤ -3
in ≥ 2 replicates
Non-toxic genes
7437
ATG9A Ratio
Z-Score (robust) ≤ -3 in all replicates
ATG9A Ratio
Z-Score (robust) ≤ -2 in all replicates &
≤ -3 in 2 replicates
ranked -> top genes selected
92
Hits
Additional Genes
58
Secondary Screen
ATG9A Ratio
Z-Score (robust) ≤ -3 in all replicates
ATG9A Ratio
Z-Score (robust) ≤ -2 in all replicates &
≤ -3 in 2 replicates
14
1° Hits
2° Hits
30
B
Primary Screen
Z–Score Robust
Secondary Screen
Z–Score Robust
−4
−6
−8
−8
−6
−4
NR3C1
RNF31
MTMR14
NTRK1
ADAM22
TMEM94
CCT6A
PDE2A
NUTF2
ADORA1
TDP1
TMEM59
NDUFB7
SEMA4D
TAZ
BTN2A1
NPM1
ELOB
KANSL2
ASIC3
NEK2
NPPA
TPCN2
NR2F1
ERBIN
HEATR5B
TNPO1
ADCYAP1
DUSP23
SMR3A
MUC21
KHK
FKBP1A
EFNA1
ANPEP
KCNQ4
AK1
MUC20
ARPC5
ATF6B
ARL5B
SNX29
ERN2
ALDH9A1
Performance
−3 in 2 | −3 in 2
−3 in 2 | −3 in 3
−3 in 3 | −3 in 2
−3 in 3 | −3 in 3

**Supplementary Figure 3: Overview of Primary and Secondary Screens and Comparison of Hit Genes**

*A)* Schematic overview of hit identification strategy for the primary and secondary screen. The ‘Synthego Human Druggable Genome’ sgRNA library targeted 8,478 genes. Toxic sgRNAs were excluded based on reduction of cell count by at least 3 standard deviations in two or more replicates, resulting in 7,437 non-toxic genes. Non-toxic genes were ranked according to ATG9A ratio reduction. A total of 92 genes reduced ATG9A ratio with a z-score robust < -3 in all replicates and were classified as hits. For secondary screening, additional 58 genes were selected from the top of the list of genes with a z-score robust < -3 in two replicates and < -2 in the third replicate, ranked by their mean ATG9A ratio across all replicates.

*B)* Comparison of confirmed hits from the secondary screen with primary screen results. Data are presented as mean z-scores robust across all replicates in the respective screen. Colors indicate if z-score robust of < -3 was achieved in only two replicates or across all three replicates.

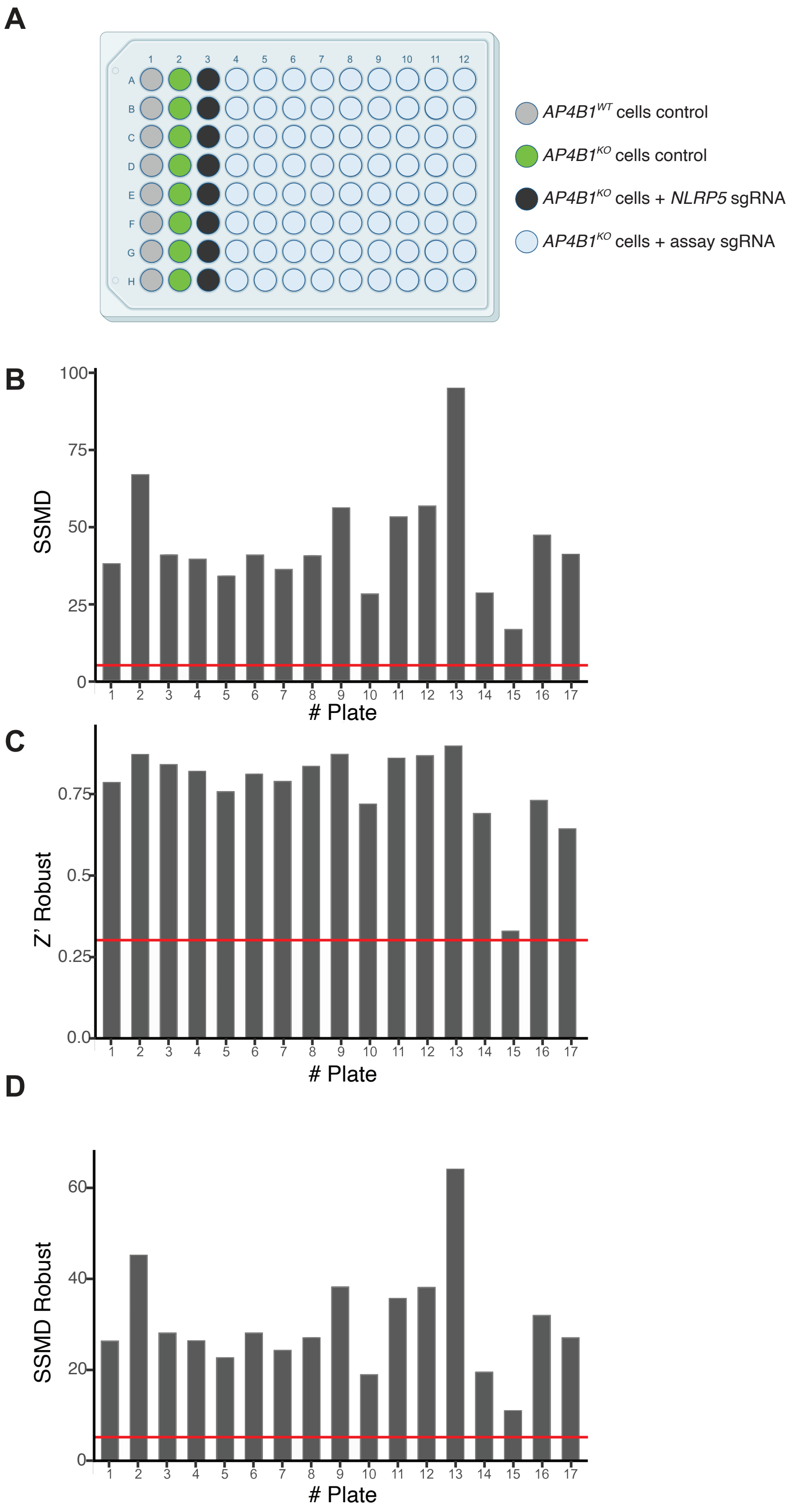
A
AP4B1^WT cells control
AP4B1^KO cells control
AP4B1^KO cells + NLRP5 sgRNA
AP4B1^KO cells + assay sgRNA
B
SSMD
### Plate
C
Z' Robust
### Plate
D
SSMD Robust
### Plate

**Supplementary Figure 4: Validation Experiments Plate Design and Quality Control Metrics**

A) Plate layout for validation experiments. Untransfected *AP4B1*$^{WT}$ cells served as positive control. Negative controls consisted of *AP4B1*$^{KO}$ cells untransfected or with transfection of sgRNAs targeting the non-essential gene *NLRP5*. This figure was created with BioRender.com.

B) Strictly standardized mean difference (SSMD) for all 17 plates of the validation experiments. Horizontal line indicates the predefined quality control threshold (SSMD > 5).

C) Z'-factor robust for all 17 plates of the validation experiments. The horizontal line indicates the predefined quality control threshold (Z' > 0.3).

D) SSMD robust for all 17 plates of the validation experiments. Horizontal line represents predefined quality control threshold (SSMD robust > 5).
